# Deep Learning-based structural and functional annotation of Pandoravirus hypothetical proteins

**DOI:** 10.1101/2023.12.02.569716

**Authors:** Joseph L. Horder, Abbie J. Connor, Amy L. Duggan, Joshua J. Hale, Frederick J. McDermott, Luke E. Norris, Sophie J.D. Whinney, Shahram Mesdaghi, David L. Murphy, Adam J. Simpkin, Luciane V. Mello, Daniel J. Rigden

## Abstract

Giant viruses, including Pandoraviruses, contain large amounts of genomic ‘dark matter’ - genes encoding proteins of unknown function. New generation, deep learning-based protein structure modelling offers new opportunities to apply structure-based function inference to these sequences, often labelled as hypothetical proteins. However, the AlphaFold Protein Structure Database, a convenient resource covering the majority of UniProt, currently lacks models for most viral proteins. Here, we apply a panoply of predictive methods to protein structure predictions representative of large clusters of hypothetical proteins shared among four Pandoraviruses. In several cases, strong functional predictions can be made. Thus, we identify a likely nucleotidyltransferase putatively involved in viral tRNA maturation that has a BTB domain presumably involved in protein-protein interactions. We further identify a cluster of membrane channel sequences presenting three paralogous families which may, as seen in other giant viruses, induce host cell membrane depolarization. And we identify homologues of calcium-activated potassium channel beta subunits and pinpoint their likely Acanthamoeba cellular alpha subunit counterparts. Despite these successes, many other clusters remain cryptic, having folds that are either too functionally promiscuous or too novel to provide strong clues as to their role. These results suggest that significant structural and functional novelty remains to be uncovered in the giant virus proteomes.

## Introduction

Since 2003 and the discovery of Mimivirus, giant viruses have put into question the traditional definition of viruses (Koonin and Yutin, 2018; Brandes and Linial, 2019). Giant viruses have been defined as being visible under a light microscope, with a recent study defining these viruses of unprecedented size as infecting eukaryotes and containing at least 500 protein-coding genes (Colson, La Scola and Raoult, 2017; Brandes and Linial, 2019). There have been further discoveries of giant viruses since Mimivirus, resulting in the classification of numerous families: *Mimiviridae, Pithoviridae, Pandoraviridae, Phycodnaviridae* and *Mollivirus (Brandes and Linial, 2019)*. Giant viruses are classified as belonging to the monophyletic group of nucleocytoplasmic large DNA viruses (NCLDVs) by phylogenetic analyses. This demonstrates the retention of most of the approximately 40 core genes that define the NCLDV group, which infect eukaryotes and reproduce within the cytoplasm of the host cell (Koonin and Yutin, 2018; Brandes and Linial, 2019). Giant virus is a literal term, with genomes reaching 2.5Mb which is considerably larger than typical NCLDVs and rivals typical bacterial genome size (Koonin and Yutin, 2018). Giant virus genomes can encode ∼1000 proteins, which is significantly more than is typical for viruses: only 0.3% of known viral genomes encode ≥500 proteins, with 69% encoding fewer than ten (Brandes and Linial, 2019). A large proportion of the products of putative protein coding regions of viral genomes remain uncharacterised (Brandes and Linial, 2019), also known as ‘dark matter’ (Koonin, Dolja and Krupovic, 2022).

The evolutionary origin of giant viruses has been the topic of debate, with the theory of a fourth domain of life being promoted and contested (Yutin, Wolf and Koonin, 2014; Colson *et al*., 2018; Koonin and Yutin, 2018; Abrahão and La Scola, 2019). There have been two evolutionary models proposed (Yutin, Wolf and Koonin, 2014; Koonin and Yutin, 2018; Brandes and Linial, 2019). First, the reductive model which favours the theory that an extinct cellular ancestor genome reduced in size, resulting in dependency on the host cell. The presence of universal cellular genes, components of translation machinery for example, within Mimivirus that appeared to not fall into any of the three domains of life supported this hypothesis. The second is the expansive model, which suggests that smaller viruses are the ancestors of giant viruses, contradicting the fourth domain hypothesis (Yutin, Wolf and Koonin, 2014; Koonin and Yutin, 2018; Brandes and Linial, 2019). A recent phylogenetic study revealed that the majority of translation-related genes of giant viruses have eukaryotic lineages and closely related eukaryotic homologues. This suggests that they have been captured from hosts, contradicting the reduction theory of a single cellular ancestor (Fischer *et al*., 2010; Koonin and Yutin, 2018; Chelkha *et al*., 2020; Sun *et al*., 2020; Belhaouari *et al*., 2022). The phylogenetic demonstration that different giant virus genes originate in any of the three domains of life supports the occurrence of exchange of genetic material between viruses and other organisms, providing further evidence for the expansion theory (Iyer *et al*., 2006; Koonin and Yutin, 2018; Brandes and Linial, 2019; Tekle *et al*., 2023). Indeed, the NCLDVs as a whole undergo extremely dynamic evolution (Brandes and Linial, 2019), again consistent with the expansion theory (Koonin and Yutin, 2018). Although studies have revealed giant viruses to belong to the NCLDVs, the evolutionary processes that have resulted in the significant genomic expansion are yet to be elucidated (Koonin and Yutin, 2018). It has been speculated that several processes are responsible, including gene duplication, horizontal gene transfer and *de novo* gene creation (Legendre *et al*., 2018; Brandes and Linial, 2019; Poirot *et al*., 2019). Furthermore, viral genes that have been captured from hosts, a process described as ‘gene gain’ (Koonin, Dolja and Krupovic, 2022) which plays a key role in the evolution of viruses (Koonin, Dolja and Krupovic, 2022). The protein function of the captured genes can either be directly used by the virus, or be repurposed through exaptation in which the structure of the acquired protein remains mostly the same, but the function is changed. This exaptation alongside *de novo* gene creation are the likely methods by which viral genomes acquire genomic dark matter (Koonin, Dolja and Krupovic, 2022).

Recently, the genomes of giant viruses have been shown to encode metabolic genes involved in energy production including glycolysis, gluconeogenesis, tricarboxylic acid cycle, photosynthesis and =-oxidation, as well as enzymes required for protein synthesis (Belhaouari *et al*., 2022). Alongside the translation-related genes mentioned above, other gene products related to RNA maturation, DNA maintenance and proteostasis have also been identified in giant viruses (Fischer *et al*., 2010; Yutin *et al*., 2013; Belhaouari *et al*., 2022). The presence of metabolic genes in giant viruses is thought to be involved in maintaining an optimal environment within the host in which the virus can replicate (Belhaouari *et al*., 2022). Furthermore these genes may provide the infected host cells with an ability to withstand harsh environments, increasing the survival of infected cells and consequently the replication of the virus (Belhaouari *et al*., 2022).

The first Pandoravirus to be discovered was *Pandoravirus salinus* in 2013 (Philippe *et al*., 2013). Pandoraviruses are among the largest of known giant viruses, with genomes exceeding 2.5Mb (Philippe *et al*., 2013; Koonin and Yutin, 2018; Legendre *et al*., 2018). For comparison, the genome of *Staphylococcus aureus* is ∼2.9Mb in size, illustrating that Pandoravirus genomes have entered the size realm of prokaryotes (Baba *et al*., 2008). Pandoravirus morphology is unique amongst giant viruses, being ovoid in shape and possessing an apical pore which facilitates the passing of particle contents into the *Acanthamoeba* host cytoplasm (Philippe *et al*., 2013; Legendre *et al*., 2018; Brahim Belhaouari *et al*., 2019; Rolland *et al*., 2019). As of June 2022 the *Pandoraviridae* family consists of 17 members: *P. pampulha, P. quercus, P. celtis, P. inopinatum, P. salinus, P. dulcis, P. massiliensis, P. neocaledonia, P. braziliensis, P. macleodensis, P. hades, P. persephone, P. japonicus, P. tropicalis, P. kadiweu, P. aubagnensis, P. belohorizontensis (Brahim Belhaouari et al., 2022)*. Pandoraviruses *salinus, inopinatum, quercus, dulcis, macleodensis* and *neocaledonia* have genome sizes ranging between ∼1.8 – 2.5Mb, encoding ∼926 – 1430 predicted Open Reading Frames (ORFs), approximately 70% of which encode uncharacterised proteins (Legendre *et al*., 2018). These Pandoraviruses have previously had their protein-coding genes functionally annotated via BLASTp (Altschul *et al*., 1990) and InterProScan (Jones *et al*., 2014; Legendre *et al*., 2018). 11.7-18.9% of the characterised proteins are deemed to have the function of protein-protein interactions (Legendre *et al*., 2018). Outside this broad category, less than 20% of predicted Pandoravirus proteins have an informative function attributed to them (Legendre *et al*., 2018).

The genes that are shared among *P. salinus, P. inopinatum, P. quercus, P. dulcis, P. macleodensis* and *P. neocaledonia* are referred to as core genes, 65% of which are unclassified, which is intriguing as these common genes are presumably essential (Legendre *et al*., 2018). Furthermore, 95% of the predicted proteins that are strain-specific were functionally unassigned (Legendre *et al*., 2018). Although Pandoraviruses contain machinery for DNA replication and transcription, mass spectrometry proteomic analysis of *P. salinus, P. dulcis, P. quercus* and *P. neocaledonia* particles presented no indication of the Pandoravirus-encoded RNA polymerase, suggesting that the transcriptional machinery residing in the host nucleus is necessary during the beginnings of infection (Legendre *et al*., 2018).

Hypothetical proteins (HPs) are gene products of the ORFs of a sequenced genome that are predicted to be expressed but lack evidence of *in vivo* existence and functional annotation (Kumar *et al*., 2014; Ijaq *et al*., 2019; Uddin *et al*., 2019). HPs are not uncommon, being found even in model organisms, and typically constitute around 30-40% of genes from a newly sequenced bacterial genome (Ashrafi *et al*., 2019). For example, 27% of the genome of *Mycobacterium tuberculosis*, the bacterium that causes tuberculosis, encodes HPs (Yang, Zeng and Tsui, 2019). HPs from numerous organisms have successfully been functionally annotated computationally through methods including structural bioinformatics approaches (da Costa *et al*., 2018; Ashrafi *et al*., 2019; Ijaq *et al*., 2019; Yang, Zeng and Tsui, 2019). The availability of AlphaFold 2 (AF2; ref) and the AlphaFold Protein Structure Database () provide new opportunities for structure based function annotation (eg) and detection of novel protein families (Bordin *et al*., 2023; Durairaj *et al*., 2023; Hernandez *et al*., 2023). Notably, viral proteins are currently largely absent from the AlphaFold Database, somewhat complicating their study: nevertheless, AF2 models have proved useful for virus phylogenetic analysis (Evseev *et al*., 2023) and function annotation (Say *et al*., 2023).

Here we apply remote homology detection and structural bioinformatics methods to predict the function of several HPs from Pandoraviruses *neocaledonia, quercus, dulcis* and *salinus.* We take advantage of the unprecedented model accuracy of AlphaFold 2 to provide the basis for structure-based function inference. We confidently annotate a number of protein families in detail, but others remain functionally cryptic despite the availability of reliable models, partly as a result of their significant structural novelty with respect to structural databases.

## Materials and Methods

### Identification of clusters of hypothetical proteins

The proteins predicted to be encoded by *P. neocaledonia, P. quercus, P. dulcis* and *P. salinus* were downloaded from UniProt (UniProt Consortium, 2023) resulting in 1081, 1179, 1070 and 1430 proteins respectively and 4760 in total. HHSuite3 (Steinegger *et al*., 2019) was used to build a custom database of profile Hidden Markov Models (HMMs) (Söding, 2005) for the 4760 total proteins. HHSearch (Söding, 2005; Steinegger *et al*., 2019) was used to detect homology between the 4760 proteins in an ‘all-against-all’ fashion locally with default parameters and 2 iterations of HHblits (Remmert *et al*., 2011) for Multiple Sequence Alignment (MSA) generation. Addition of secondary structure via the addss.pl script from HHSuite3 was not applied at this step. The resulting probability scores were processed into a matrix using locally written scripts (adapted from Mesdaghi, personal communication) which removed any hits a protein had against itself. Where there were multiple hits between two proteins, only the top hit was retained. CLANS (Frickey and Lupas, 2004) was then employed to use this matrix to organise the 4760 proteins into network-based clusters of homologous sequences. A threshold was set meaning that only clusters with at least 10 members were output. Clusters containing only hypothetical (uncharacterized) proteins were taken forward for further analysis.

### Sequence-based predictive methods

Clusters were aligned using MAFFT (Katoh and Standley, 2013) and Muscle (Edgar, 2004), which typically gave similar results, and visualised with Jalview (Procter *et al*., 2021). The alignment was used to select one, or occasionally two, representatives for sequence analysis and modelling. Representatives were selected to be typical of the overall set of sequences and, in particular, to contain all aligned sequence blocks.

Representative sequences (whose UniProt codes are shown in Table 1) were submitted to DeepTMHMM (Hallgren *et al*., 2022) for prediction of transmembrane alpha-helices, SignalP 6.0 (Teufel *et al*., 2022) for signal peptide prediction, DeepCoil (Ludwiczak *et al*., 2019) for coiled-coil protein architecture, DeepLoc 2.0 (Thumuluri *et al*., 2022) for predicted subcellular localisation and fIDPnn (Hu *et al*., 2021) for prediction of intrinsically disordered regions. IUPred3 with ANCHOR2 (Erdős, Pajkos and Dosztányi, 2021) was used for disordered binding site predictions and Phobius (Käll, Krogh and Sonnhammer, 2007) for predictions for transmembrane topology. (PSI-)BLAST (Altschul *et al*., 1997) and programs of the hmmer suite (Eddy, 2009) were used to identify any homologues outside pandoraviruses and HHsearch (Söding, 2005) used for the detection of homology between cluster representative sequences and Pfam families (Mistry *et al*., 2021) or structurally characterised proteins in the PDB (wwPDB consortium, 2019). Where necessary, BLAST (Altschul *et al*., 1997) was also used to identify orthologues by reciprocal BLASTing.

**Table 1:** Characteristics and sequence-based predictions for the clusters of Pandoravirus hypothetical proteins.

| Paper cluster number | Number of members and in individual viruses (Pn, Pq, Pd, Ps) | Selected representative (length) | Signal peptide detected by SignalP-6.0? | Transmembrane helices detected by DeepTMHMM | Localization(s) predicted by DeepLoc 2.0 | Major (>20 residues) regions of predicted intrinsic disorder with fIDPnn | Gene Ontology terms predicted for Molecular Function by deepFRI. * indicates that only the highest-scoring and most specific predictions are shown |
| --- | --- | --- | --- | --- | --- | --- | --- |
| 1 | 58 (5, 20, 15, 18) | A0A2U7UAA1 (289) | No | No | Cytoplasm |  |  |
| 2 | 36 (8, 8, 9, 11) | A0A2U7U7X3 (184) | No | Yes<br>TM1 15-35<br>TM2 153-175 | Cell membrane |  | GO:0009055 electron transfer activity (0.61)<br>GO:0016491 oxidoreductase activity (0.52) |
| 3 | 30 (8, 4, 8, 10) | A0A2U7UDC4 (296) | No | No | Cytoplasm, nucleus |  | GO:0043167 ion binding (0.58) |
| 4 | 27 (5, 9, 7, 6) | A0A2U7U8Z7 (228) | Yes residues 1-31 | No | Extracellular |  |  |
| 5 | 26 (6, 6, 8, 6) | A0A2U7UDI8 (577) | No | Yes<br>TM1 62-82<br>TM2 101-119<br>TM3 135-154<br>TM4 161-181<br>TM5 200-215<br>TM6 237-257<br>TM7 316-335<br>TM8 354-374<br>TM9 380-401<br>TM10 410-425<br>TM11 448-469<br>TM12 483-503 | Lysosome/vacuole | 525-553 | GO:0022857 transmembrane transporter activity (0.97)<br>GO:0015291 secondary active transmembrane transporter activity (0.56) * |
| 6 | 25 (7, 6, 5, 6) | A0A2U7UCR8 (691) | No | No | Cytoplasm |  |  |
| 7 | 23 (6, 4, 4, 9) | A0A2U7UAG2 (912) | No | No | Cytoplasm, Nucleus | 1-38, 172-196, 577-641 | GO:0043167 ion binding (0.51) |
| 8 | 22 (3, 8, 6, 5) | S4VVH3 (275) | No | No | Mitochondrion |  | GO:0016829 lyase activity (0.68)<br>GO:0016835: carbon-oxygen lyase activity (0.61) |
| 9 | 21 (3, 7, 5, 6) | A0A2U7U902<br>(283) | No | No | Cytoplasm |  | GO:0016787 hydrolase activity<br>GO:0046872 metal ion binding |
| 10 | 21 (5, 6, 4, 6) nqds | S4VZQ1 (234) | No | Yes<br>TM1 43-63<br>TM2 71-93 | Endoplasmic reticulum |  | GO:0015318 inorganic molecular entity<br>transmembrane transporter activity (0.79)<br>GO:0015077 monovalent inorganic cation<br>transmembrane transporter activity (0.51) * |
| 11 | 16 (4, 4, 4, 4) | S4VZY3 (441) | No | No | Cytoplasm |  | GO:0043167 ion binding (0.88)<br>GO:0005524 ATP binding (0.71) * |
| 12 | 16 (4, 4, 4, 4) | S4VS46(184) | Yes, residues<br>1-18 | Yes<br>TM1 89-111 | Lysosome/vacuole | 133-182 |  |
| 13 | 13 (2, 3, 5, 3) | S4VZF8 (182) | No | Yes,<br>TM1 55-73<br>TM2 158-177. | Endoplasmic reticulum | 1-39 |  |
| 14 | 12 (0, 4, 4, 4) | A0A2U7U965<br>(116) | No | No | Extracellular | 1-20 |  |

### Phylogenetics

MEGA (Hall, 2013) was used for phylogenetic analysis using protein sequence alignments generated with MAFFT (Katoh and Standley, 2013). Neighbour-joining and Maximum Likelihood (ML) methods were employed for tree construction with bootstrapping used to assign confidence to nodes. As results were generally similar, only the ML trees were used for further analysis.

### Protein modelling

The ColabFold (Mirdita *et al*., 2022) webpages were used for generation of AlphaFold 2 models using default settings unless otherwise stated. The same pages were used for construction of dimer models where appropriate. In some cases, local AlphaFold 2 (Jumper *et al*., 2021) was also run. The ColabFold webpage was used for generation of OmegaFold (Wu *et al*., 2022) models using default settings unless otherwise stated. Any overall confidence scores for OmegaFold models presented here was calculated by obtaining the mean of the confidence scores of all individual residues in the model. RosETTAFold (Baek *et al*., 2021) models were generated using the Robetta Server (Kim, Chivian and Baker, 2004), using default settings.

### Structure-based function annotation

Structural neighbours were sought with DALI (‘Protein Structure Comparison by Alignment of Distance Matrices’, 1993) and FoldSeek (van Kempen *et al*., 2023). Functional sites were sought using a variety of approaches. Pockets were characterised using the CastP 3.0 server (Tian *et al*., 2018), electrostatic analysis done initially using PyMOL’s inbuilt facility with APBS (Jurrus *et al*., 2018) follow-up where appropriate, sequence conservation mapping with ConSurf (Ashkenazy *et al*., 2016) (where the number of homologues in the databases was sufficient), prediction of binding sites using surface triplets with STP (Mehio *et al*., 2010). For cluster 7, ConSurf was run providing the results of a hmmer (hmmer.org) search against the nr90_29_Apr database, applying an e-value threshold of 0.001 and aligning the results with MAFFT (Katoh and Standley, 2013). Deep learning-based function prediction was done with DeepFRI (Gligorijević *et al*., 2021) and PeSTo (Krapp *et al*., 2023), and potential protein interfaces identified using CSM-Potential (Rodrigues and Ascher, 2022). SWORD2 (Cretin *et al*., 2022) was used to define domain boundaries and ATPbind used to predict the nucleotide-binding site of cluster 7 (Hu *et al*., 2018).

## Results

### Identification of clusters entirely comprising hypothetical proteins

The clustering exercise among the proteomes of four Pandoraviruses described in Methods identified 70 clusters containing 10 or more sequences. Twenty-three of these consisted entirely of hypothetical proteins of which 14 (see Table 1) were taken forward.

The clusters ranged in size from 58 members (cluster 1) to 12 members (cluster 14). In all except cluster 14 there were multiple representatives from each of the four pandoraviruses, *P. neocaledonia, P. quercus, P. dulcis* and *P. salinus*. Although not all cluster members were full-length and highly conserved positions were sometimes absent in some cluster members, these cluster sizes can reasonably be understood as reflecting important functions encoded by the proteins found therein. The largest number of homologues in a single virus was 20 for *P. quercus* and cluster 1. Cluster 14 contained no sequences from *P. neocaledonia*, but four from each of the other viruses. None of the representative sequences contained coiled-coil predictions of longer than 10 residues but five contained at least one predicted transmembrane helix and five contained at least one major region of predicted intrinsic disorder longer than 20 residues. Predicted subcellular localizations were diverse (Table 1) and were predicted on the basis that the viral proteins would be processed by the host machinery. The representative sequences of these were subjected to detailed analysis and structure-based annotation using AlphaFold 2 (AF2) models.

### Confident functional assignments from AlphaFold 2 modelling

#### Cluster 7 contains nucleotidyltransferase enzymes

Cluster 7 consisted of 23 sequences of HPs from *P. quercus*, *P. neocaledonia*, *P. dulcis* and *P. salinus* (4, 6, 4 and 9 sequences respectively) with A0A2U7UAG2 of *P. quercus* being selected as the representative protein with a length of 912AA. The full sequence of A0A2U7UAG2 was uploaded to the HHPred server against the PDB_mmCIF70 and Pfam-A_v35 databases with default parameters. The results fell within 2 clear regions of the sequence (Table 1). Region 1 (residues ∼50-165) returned hits against BTB domain-containing structures, with the top PDB hit against Kelch-like ECH-associated protein BTB domain (6I2M_A, Prob = 92.21%) and top Pfam hit of BTB domain (PF11822.11, Prob = 90.07%) with almost full alignments. Region 2 (residues ∼395-580) returned a top characterised PDB hit against Poly A polymerase/RNA nucleotidyltransferase (6QY6_A, Prob = 95.48%), with only partial alignment against the hit. The Pfam results identified Poly A polymerase head domain (PF01742.23, Prob = 95.07%) and Nucleotidyltransferase domain (PF06042.14, Prob = 90.03%) with essentially full alignments against the hits. The IUPred3 (Erdős, Pajkos and Dosztányi, 2021) server predicted two putative globular domains between residues 52-560 and 653-912. Interestingly, the BTB and PolyA Polymerase head/nucleotidyltransferase domains highlighted by HHPred fell within globular domain 1 while the HHPred search did not produce any results for the second predicted globular domain.

A model of A0A2U7UAG2 (Fig 1a) was predicted with AF2 and trimmed by removing residues with a pLDDT score <70 to leave only the most confidently modelled regions (Fig 1b). The DALI server was used to identify structural similarities between the trimmed model and known structures in PDB25 (Table 2). In agreement with the HHPred results, the DALI structural search results identified confident hits against tRNA processing enzyme complex 2 (3wfr-H, Z score = 10.5), Poly(A) polymerase (3aqk-A, Z score = 8.1) and BTB/POZ domain-containing protein (2vkp-B, Z score = 7.7). Further structure based searches against the AFDB50 and MGNIFY_ESM30 using the FoldSeek server identified further confident structural matches (Table 2) but only BTβ-domain containing proteins were functionally annotated with other regions corresponding to hypothetical or uncharacterised proteins. The FoldSeek search against the PDB identified both BTB domain containing proteins (top hit 5daf-A, Prob=1.00, TM score=0.55) and tRNA processing enzymes (top hit 3wfp-G, Prob=0.98, TM score=0.55). Restricting the FoldSeek search to viruses returned 0 results against the AFDB50, but it is known that virus proteins are largely absent from the current AFDB. One viral protein match was identified in the PDB, a crystal structure of vaccinia virus protein A55 BTB-Back domain in complex with human Cullin-3 N-terminus (6i2m_A, Prob=0.97, TM score=0.49). Indeed, 6i2m_A was the top BTβ-domain containing protein hit identified in the HHPred search described above.

**Figure 1.**
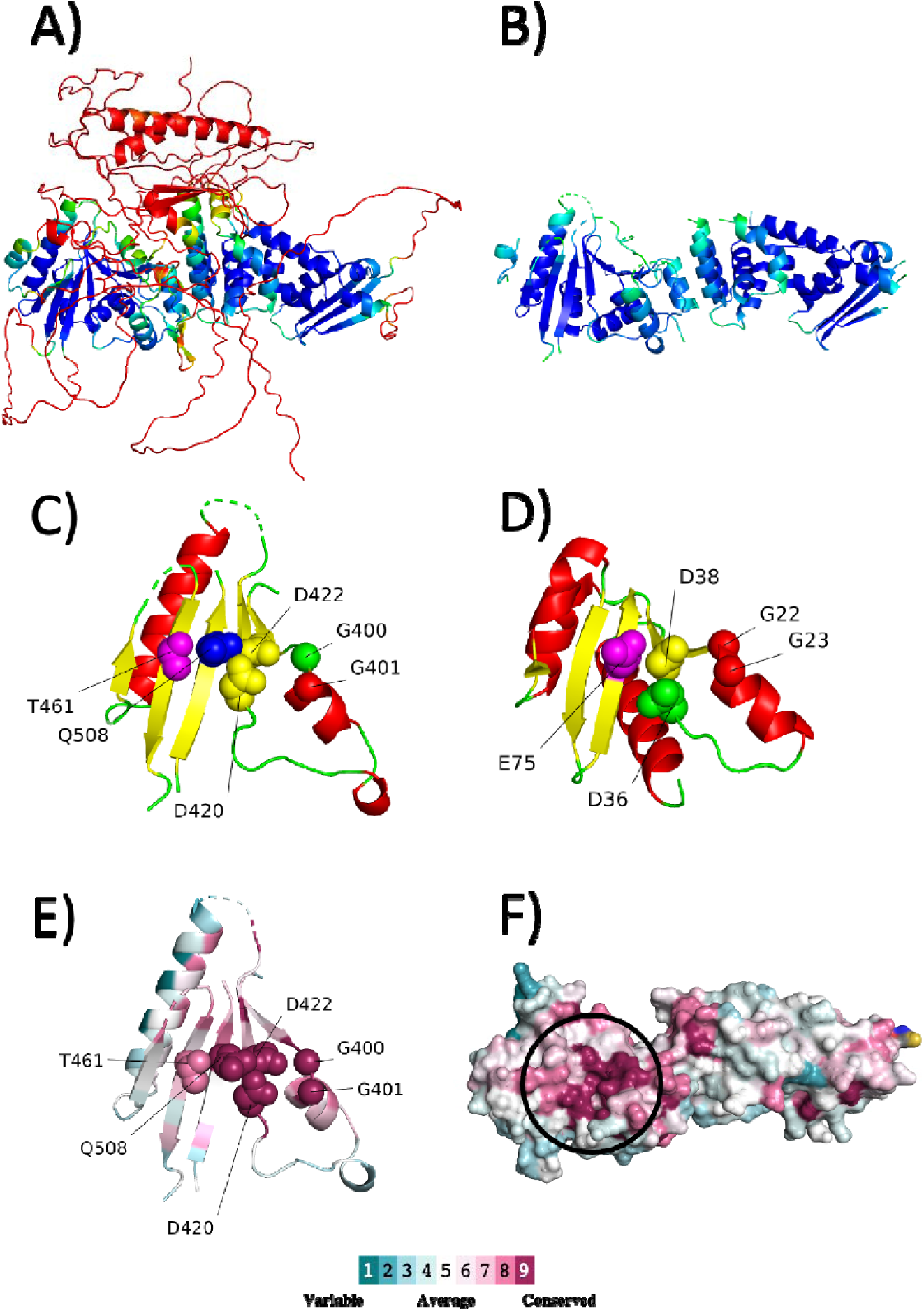
A. The full model of A0A2U7UAG2 coloured by pLDDT score as specified by ColabFold. B. The trimmed model of A0A2U7UAG2 to include only residues with a pLDDT score ≥ 70. C. The Polβ superfamily consensus fold and motif present within the model of A0A2U7UAG2. The structure is coloured by secondary structure with the motif residues side chains labelled and displayed as spheres. The residue aligning with the third carboxylate of 6QY6_A (T461) via HHPred is coloured in magenta. The residue that structurally overlays with PyMOL cealign with the third carboxylate of 6QY6_A (Q508) is coloured in blue. D. The Polβ superfamily consensus fold and motif present within the structure of 6QY6_A. The structure is coloured by secondary structure with the motif residues side chains labelled and displayed as spheres. The third carboxylate is coloured in magenta. E. Sequence conservation mapping by ConSurf to the Polβ superfamily consensus fold and motif present within the model of A0A2U7UAG2. The mapping is coloured as a spectrum, which colour to conservation shown in the figure. The Polβ motif residues as described in C are labelled and shown as side chains. F. Sequence conservation mapping by ConSurf to the pLDDT trimmed model of A0A2U7UAG2. The site of the predicted Polβ superfamily consensus fold and motif is circled.

**Table 2:**
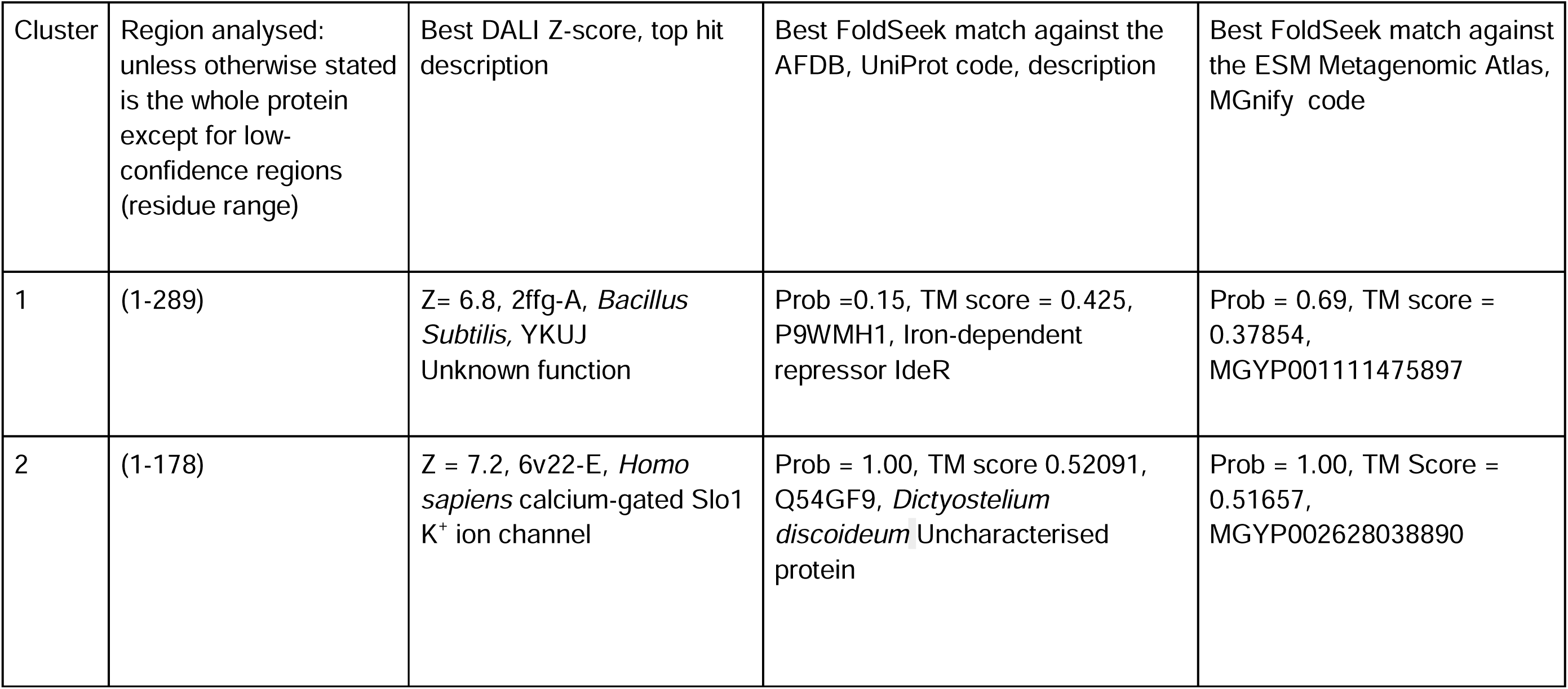
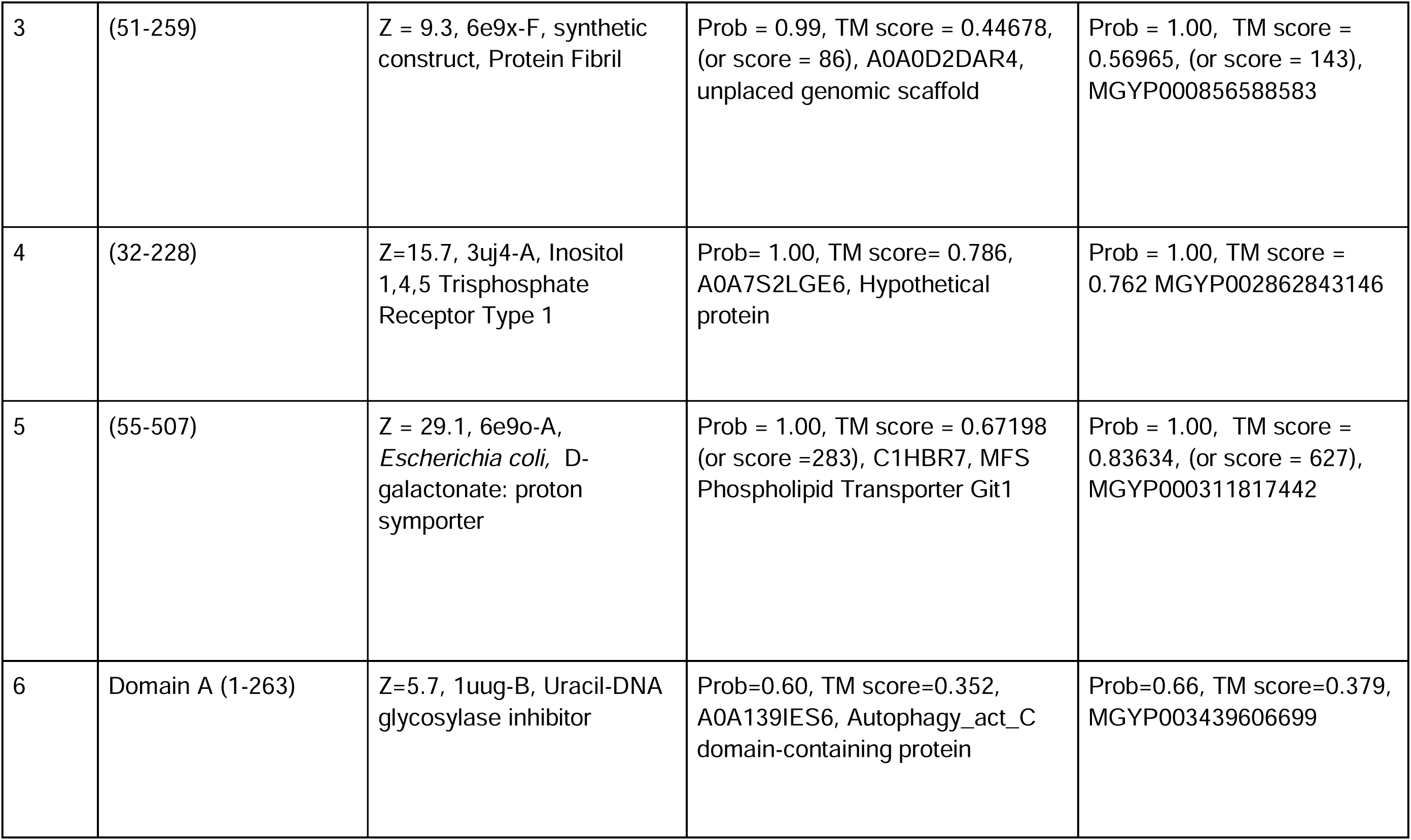
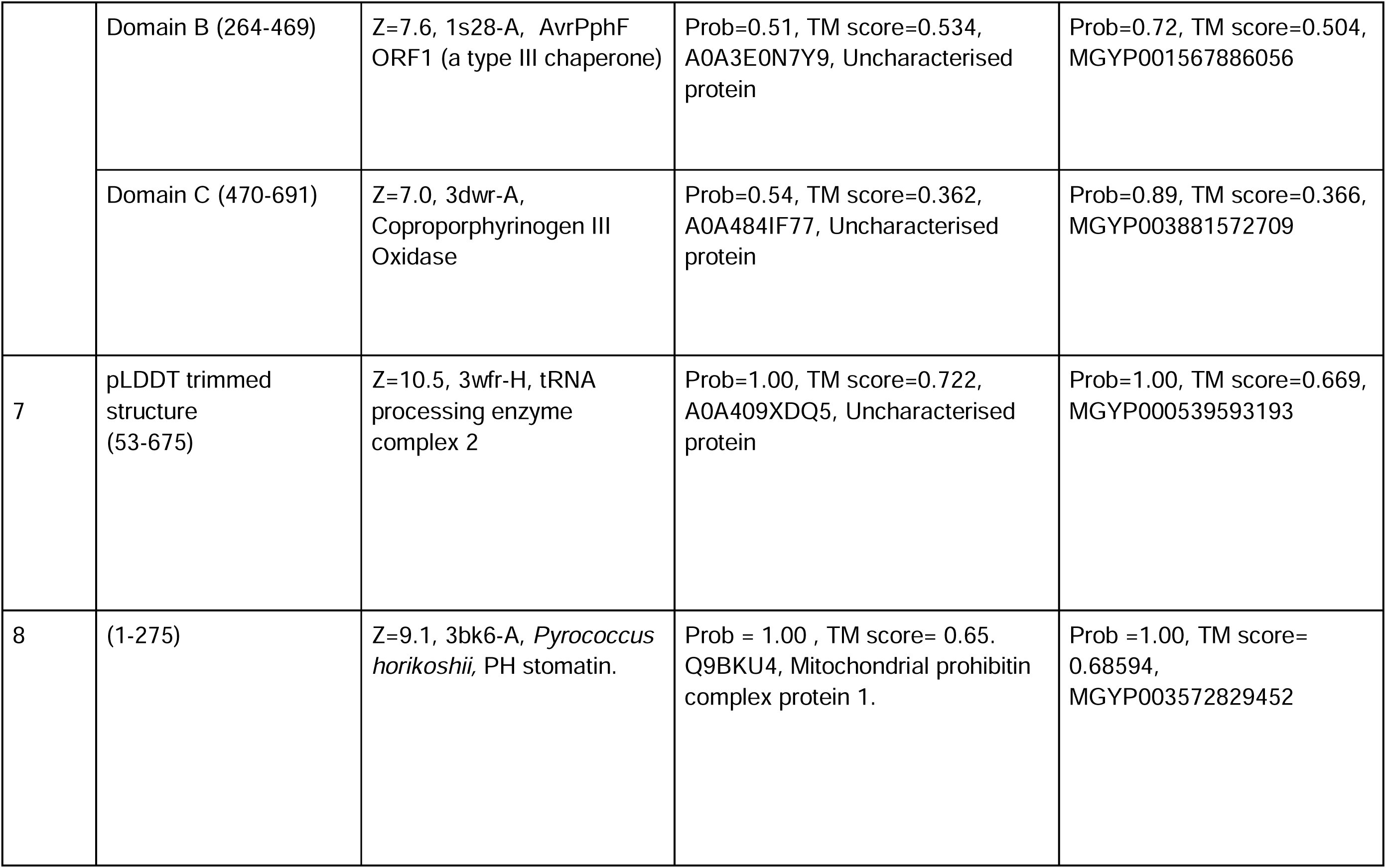
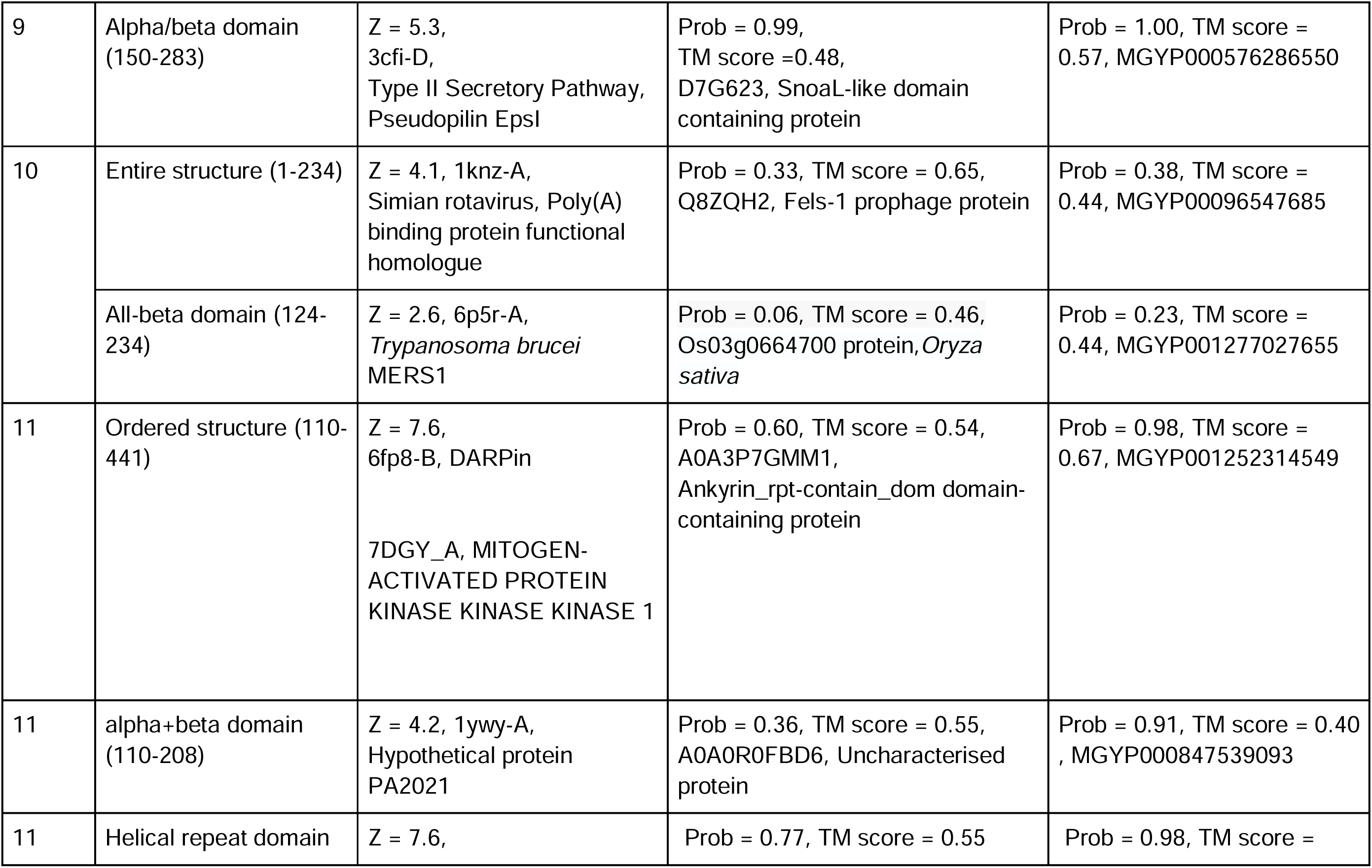
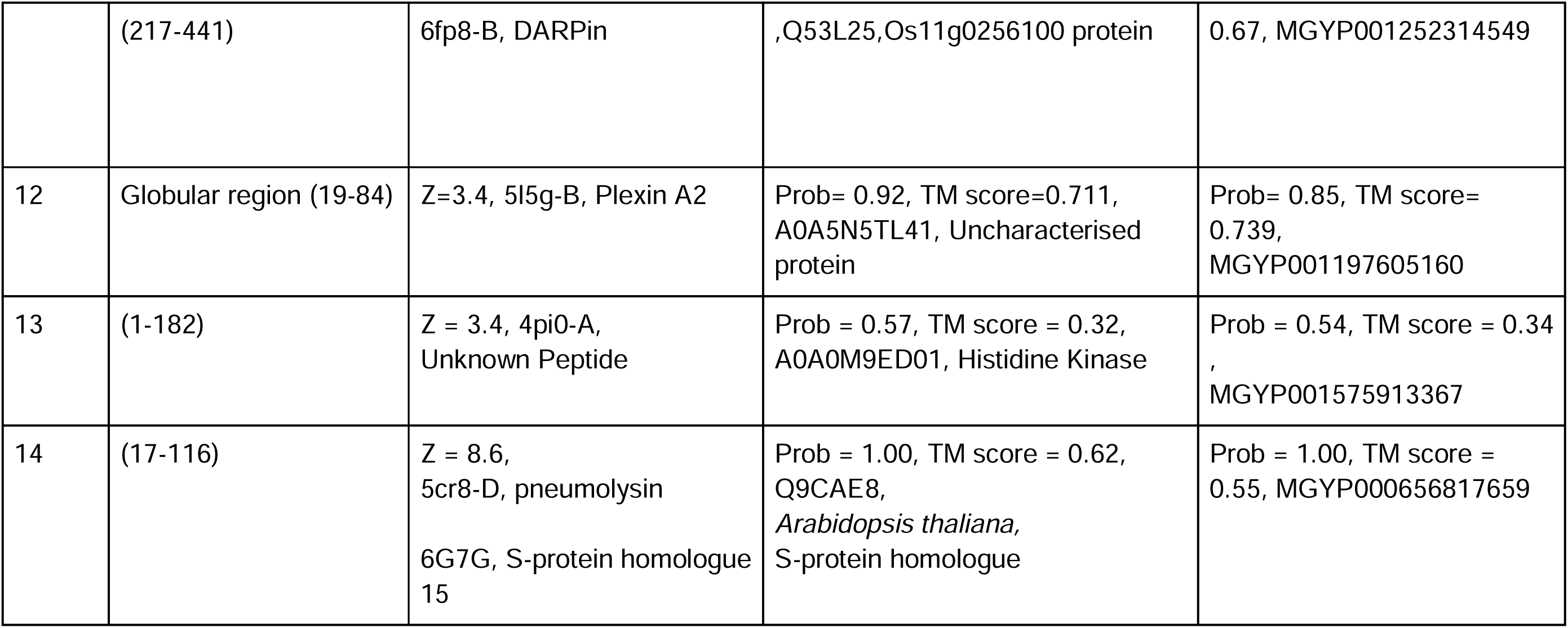
Matches to AF2 models of cluster representatives in structure databases.

#### The cluster 7 nucleotidyltransferase domain

Class II nucleotidyltransferases, including both PAPs and CCA-adding enzymes (CAEs), catalyse the addition of nucleotides to RNA and belong to the Polymerase β superfamily (Polβ) whose substrates are ATP and mRNA for PAPs and ATP, CTP and tRNA for CAEs (Balbo and Bohm, 2007; Vörtler and Mörl, 2010; Yang *et al*., 2013; Tomita and Yamashita, 2014). The Polβ family is defined by the presence of a signature fold and motif that coordinates two metal ions to facilitate catalysis (Betat, Rammelt and Mörl, 2010). The fold consists of two α-helices either side of a five stranded antiparallel β-sheet (Betat, Rammelt and Mörl, 2010). Both of these class II nucleotidyltransferases (NTs) share a homologous region in which the catalytic site is located (Betat, Rammelt and Mörl, 2010; Toh *et al*., 2011). Several conserved motifs essential for function are present in the NT region: the Polβ motif hG(G/S)X(9-13)Dh(D/E)h (motif A), RRD (motif B) and EDXXR (motif C) where h denotes a hydrophobic residue and X any residue (Just *et al*., 2008; Betat, Rammelt and Mörl, 2010; Toh *et al*., 2011). Class I and II nucleotidyltransferases are also separated based on their phylogeny. Class I PAPs are found in eukaryotes and class I CAEs in archaea. Class II PAPs on the other hand are found in bacteria and class II CAEs are both bacterial and eukaryotic (Betat, Rammelt and Mörl, 2010).

The model of A0A2U7UAG2 displays the Polβ fold described above. Significantly, a comparison of this fold between the model and the top HHPred hit against the PDB of known function (6qy6-A) reveals the Polβ signature motif (motif A) to be present in the correct structural location of the modelled fold, satisfying the criterion to be considered a member of the superfamily and therefore adding confidence to the HHpred and DALI results against class II nucleotidyltransferases. Consurf results (Fig 1) showed the signature Polβ motif and the active site to be conserved. Alignment of all sequences in cluster 7 (bar 4 short sequences) with MAFFT showed conservation of the Polβ catalytic motifs, suggesting that a preliminary prediction of NTase function can be applied to the whole of cluster 7.

Along with the two carboxylates present in the Polβ motif (motif A), a third D/E is present in the superfamily fold that aids in the coordination of metal ions in the active site of class II nucleotidyltransferases for catalysis, as described in the original publication for the HHPred target 1miw-A (HHpred Prob > 90%) (Li *et al*., 2002; Toh *et al*., 2011; Wolkowicz and Cook, 2012). This third carboxylate is not present in the model of A0A2U7UAG2. However, the HHPred alignment of A0A2U7UAG2 aligns a conserved threonine (T461) to the third carboxylate of 1miw-A. Furthermore, a highly conserved glutamine (Q508) resides in a region corresponding to the third carboxylate of 1miw-A and PeSTo webserver results support the suggestion that it, along with D420 and D22, comprise an ion binding site (Fig 1). Finally, whilst this third carboxylate is present in the HHPred NTase hits, it has been demonstrated that class II, unlike class I, CAEs require only the two carboxylates present in motif A to maintain function (Hou *et al*., 2005), suggesting that a loss of the third carboxylate is not detrimental to function and further supporting the HHpred results favouring class II nucleotidyltransferase activity over class I.

Motifs B and C of class II nucleotidyltransferases described above did not appear to be present in A0A2U7UAG2. Motif B, RRD in the crystal structures of NTases, is replaced by DFD in A0A2U7UAG2, although the conservation of residues F524 and D525 is suggestive of functional importance. Motif C is not recognisably present, although it is interesting to note that two Arg residues just downstream of it, R565 and R568, are highly conserved and match residues that interact with the nucleobase in the functional pocket of 6qy6-A (de Wijn *et al*., 2021). To further investigate the putative ATP binding capabilities of A0A2U7UAG2 the ATPbind webserver (Hu *et al*., 2018) was used. Indeed, the residues predicted to bind ATP were G401, D420, D523, F524, D525, A529 and W542 in the predicted active site. G401 and D420 are within motif A whilst conserved F524 and D525 align with motif B. R565 and R568 were not identified by ATPbind but, as mentioned, their conservation and alignment with binding residues in the crystal structure (Fig 2) adds strength to the prediction of NTase activity.

**Figure 2.**
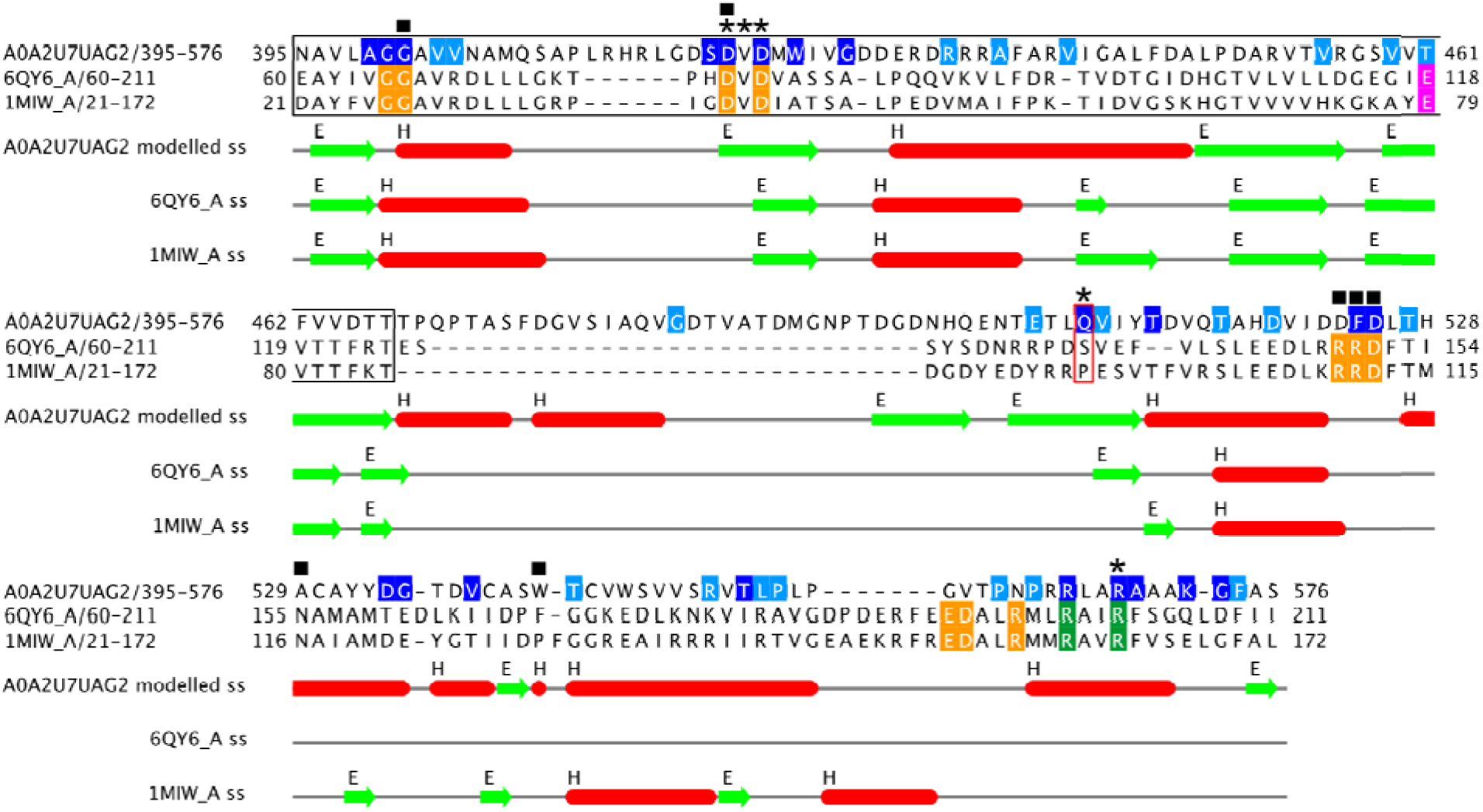
HHpred alignment of cluster 7 representative A0A2U7UAG2 with two NTase enzymes of known structure. Orange residues are motifs A - C mentioned in the text, magenta residues are the third carboxylates also mentioned in the text and dark green residues two further key functional positions. Residues in A0A2U7UAG2 are coloured according to conservation (dark blue being most conserved). Squares are used for residues predicted to bind nucleotide at the ATPbind server, asterisks for PeSTo ion binding positions. The black box marks the core Polβ superfamily consensus fold and the red box marks the putative third metal-binding residue in A0A2U7UAG2, as mentioned in the text. Secondary structure in the model and crystal structures is marked below the alignment. The alignment was prepared using Jalview (Waterhouse *et al*., 2009).

As discussed, class II PAPs and CAEs share homologous N-terminal catalytic regions (Betat et al., 2010) which has prompted research into the mechanism of specificity of ATP in PAPs as opposed to CTP and ATP in CAEs (Toh *et al*., 2011). Indeed, it is the C-terminal (CT) half of the enzymes which confers nucleotide specificity (Toh *et al*., 2011). Topologies and folds of the CT half are comparable between the two enzymes, composed entirely of helices, but with subtle differences in the organisation of the helices (Toh *et al*., 2011). A key structural feature that differs between class II PAPs and CAEs is a helix within a hand domain (Toh *et al*., 2011). In PAPs, the hand domain interacts with the catalytic pocket and likely maintains the structure of the nucleotide binding site, resulting in specificity for ATP (Toh *et al*., 2011). The same helix is present in CAEs but, rather than interacting with the pocket, it interacts with other helices in the CT (Toh *et al*., 2011), which has been speculated to increase the flexibility of the nucleotide binding pocket allowing both CTP and ATP to enter (Toh *et al*., 2011). In this case, a hand domain is not evident in the model, but the original publication of 6qy6-A (de Wijn *et al*., 2021), the top known PDB hit from HHPred, describes the EDXXR motif (motif C) as containing the base-recognizing residues of the molecule. As A0A2U7UAG2 appears to be missing this clear motif we can speculate that A0A2U7UAG2 does not retain base specificity.

#### The cluster 7 BTB domain

The BTB domain is a conserved motif for protein-protein interactions (PPIs), both with other BTB domains and non-BTB proteins, that has been implicated in several functions including protein ubiquitination and degradation (Stogios *et al*., 2005; Orosa *et al*., 2017; Han *et al*., 2019). BTB domain-containing proteins have been categorised into families depending on their specific function and can be found in conjunction with other domains, e.g. the BACK domain (Stogios and Privé, 2004; Stogios *et al*., 2005; Orosa *et al*., 2017; Han *et al*., 2019). Despite the lack of sequence similarity between the BTB domain families, central to all is a core fold with a conserved tertiary structure (Stogios *et al*., 2005), suggesting that identification of this fold is a strong indication of PPI capability. In addition, the CSM-Potential webserver (Rodrigues and Ascher, 2022) predicts the putative BTB domain containing region to be a site of PPI. Browsing the Pfam (Mistry *et al*., 2021) entry for the BTB/POZ domain (PF00651) reveals that the BTB domain has previously been annotated in Pandoravirus proteins.

The BACK domain, so named as it is often present in proteins containing BTB And C-terminal Kelch repeats (Stogios and Privé, 2004), has been identified to immediately follow the BTB domain in protein structure and is speculated to play a role in Cul3-mediated protein degradation (Stogios and Privé, 2004). Although the majority of BACK domain containing proteins have a domain architecture of BTB-BACK-Kelch (BBK) (Stogios and Privé, 2004), a BTB-BACK domain containing protein has been identified without evidence of a CT Kelch that mediates proteasomal degradation (Stogios and Privé, 2004; Orosa *et al*., 2017).

To see if the finer molecular determinants of PPI were present, we compared the A0A2U7UAG2 BTB domain with that found in 6i2m-A (containing BTB and BACK domains), the structure of which strongly binds Cul3, mediated by the BTB domain (Gao *et al*., 2019). The BTB-Cul3 interaction in 6i2m-A is driven by a conserved motif, hX(D/E) where h is a hydrophobic residue and X is any residue (Gao *et al*., 2019). The residues forming this motif in 6i2m-A are F54 (h), I55 (X) and D56 (Gao *et al*., 2019) and align with F92, A93 and E94 in A0A2U7UAG2. The functionally important F92 and E94 are highly conserved, in line with a role in a similar PPI.

#### A possible third folded domain in cluster 7

The predicted second globular region from residues 653-912 was investigated further. HHpred search of residues 653-912 identified two targets covering ∼50% of the query length; 4klk-A (probability=44.01) and 5odk-B (probability=42.66), both of which contain DUF2815, a phage-related family of proteins with unknown function. Whilst these scores are low, it is interesting to note that 5odk-B is defined as a single-stranded DNA-binding protein, a function which fits within the context of the NTase abilities discussed above. Furthermore, electrostatics analysis (not shown) showed a pronounced positive charge, consistent with a nucleic acid-binding role. However, arguing against a key functional role, the domain is much less conserved than other parts of the protein.

#### Potential in vivo role of cluster 7

The takeover of host processes during viral infection is known to be a key component of viral fitness (Gustin *et al*., 2011). Modification and exploitation of the ubiquitin system is utilised by nearly all classes of virus to interfere with the host throughout the viral life cycle, from viral entry to release, with ubiquitin-modifying machinery being present in numerous large DNA viruses (Gustin *et al*., 2011). Pandoraviruses are large DNA viruses (Brandes and Linial, 2019) and components of ubiquitin-dependent degradation pathways have accordingly been identified in *P. salinus*, likely present to disrupt host processes (Philippe *et al*., 2013). A presumed method of host ubiquitin manipulation by *Poxviridae*, a member of the NCLDV group which includes Pandoraviruses, is through BTB-BACK-Kelch proteins via interaction with Cul3 ligase complexes (Gustin *et al*., 2011). Indeed, analysis of A0A2U7UAG2 identified a likely BTB-BACK domain present. Furthermore, the top HHPred hit was against 6I2M_A, protein A55 from vaccinia virus, a poxvirus. A recent study of A55 (Gao *et al*., 2019) suggests that interaction with Cul3 may lead to degradation of previously untargeted host proteins and/or liberate proteins from degradation that are usually targeted during infection (Gao *et al*., 2019). It has also been reported that A55 may influence the immune response of the host to the virus (Beard, Froggatt and Smith, 2006).

CAEs are essential in translation, adding a CCA triplet to the 3’ end of tRNA, which brings amino acids to mRNA at the ribosome during protein synthesis, in order to reach fully functional tRNA maturity and plays a role in maintenance and repair of tRNA molecules (Vörtler and Mörl, 2010; Lyons, Fay and Ivanov, 2018; Torres *et al*., 2019; Wellner *et al*., 2019). *P. salinus* lacks components of translation machinery, such as ribosome proteins (Philippe *et al*., 2013), suggesting that host translation apparatus is required during Pandoravirus replication. However, translation-related genes are present in *P. salinus* including amino acid tRNA ligases, a SUA5-like tRNA modification enzyme and an eIF4E translation initiation factor (Park, Schimmel and Kim, 2008; Philippe *et al*., 2013). Three tRNAs for proline, methionine and tryptophan, are also present (Philippe *et al*., 2013). *P. neocaledonia* also encodes three tRNAs (Legendre *et al*., 2018), one for proline and two for methionine (NCBI (Sayers *et al*., 2022) Reference Sequence NC_037666.1). The presence of tRNAs and other translation related genes in Pandoraviruses supports the putative presence of a CAE in *P. quercus* which may be required to bring the tRNAs to full maturity for viral protein translation. Indeed, the identification of putative homologues of A0A2U7UAG2 in Pandoraviruses *quercus, neocaledonia, dulcis* and *salinus* emphasises the likely functional importance of A0A2U7UAG2 which would be associated with a CAE (Wellner *et al*., 2019).

From the analyses of A0A2U7UAG2 we can speculate that cluster 7 proteins may aid in *P. quercus* infection of *Acanthamoeba* through degradation/evasion of degradation of host proteins, possibly interfering with host inactivation of translation machinery upon viral infection (Walsh, Mathews and Mohr, 2013). We can also hypothesise that A0A2U7UAG2 matures tRNAs encoded by *P. quercus*, resulting in functional tRNAs contributing to viral protein translation.

### Cluster 2 contains homologues of calcium-activated potassium channel beta subunits

There were 36 proteins contained within cluster 876 which were represented by protein A0A2U7U7X3 with 184 residues. Sequence analysis (Table 1) suggested that the protein contained two transmembrane helices and resided in the cell membrane. Homologues are found in the other giant viruses *Mollivirus sibericum* and *Mollivirus kamchatka*. HHpred analysis gave strongly significant, >99% probability hits against Pfam (Calcium activated potassium channel beta subunit domain (PF03185.18); probability 99%) and the PDB (human calcium-activated potassium channel subunit beta-4 (PDB code 6v22)).

The AF2 prediction for the representative sequence A0A2U7U7X3 contained two medium confidence α-helices, coinciding with the transmembrane helix predictions from DeepTMHMM, connected by a highly confident β-fold domain (Fig 3). Offering strong support to the modelling, conserved residues C52 and C129 of cluster 2 are well-positioned to form a disulphide bond. In agreement with the HHpred results, a DALI search in the PDB with the prediction produced a top structural alignment against the 6v22-E beta subunit (Z score = 7.1). Examination of the structure superposition of the model on the complete channel structure in PyMOL revealed the conservation of residues for interaction with hydrophobic lipid tails - F2033 and F2037 in 6v22 correspond to F39 and F43 in the cluster 2 representative.

**Figure 3.**
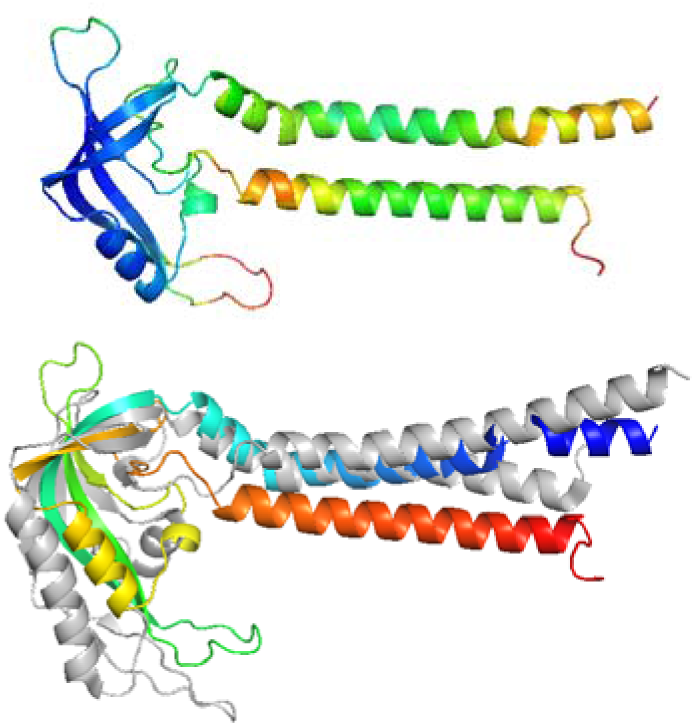
a) Cartoon of the AF2 model of the cluster 2 representative sequence, colour coded according to pLDDT confidence values from blue (high) to red (low). b) DALI superposition of the AF2 model (coloured in a spectrum from blue N-terminus to red C-terminus) with the beta subunit (chain E) of calcium-activated potassium channel beta subunit (grey PDB 6v22).

The PDB entry 6v22 is the structure of a human calcium-gated Slo1 K^+^ ion channel composed of a large alpha subunit (1065 residues) surrounded by 4 repeated auxiliary beta subunits (219 residues), the latter shown above to be homologous to A0A2U7U7X3. The beta subunits in the Slo1 channel significantly alter the biophysical properties of the ubiquitously expressed alpha-subunit, such as in changing its activation/deactivation and sensitivity to calcium (Tao and MacKinnon, 2019). These auxiliary subunits therefore endow the channel with functional diversity that enables tissue-specificity primarily associated with higher organisms (Tao and MacKinnon, 2019).

With cluster 2 seeming to comprise auxiliary beta subunit proteins, we could propose a viral function involving the interaction and interference with host ion channels to promote infection. A precedent for megavirus interference in host transmembrane ion transport is seen in giant chlorella viruses, where the virus encodes potassium ion channels inserted into host plasma membrane initiating depolarisation in order to make the release of viral DNA more favourable (Frohns *et al*., 2006). A search was therefore carried out for alpha channel subunits with which cluster 2 beta subunits might associate.

The search for alpha subunits was carried out in both Pandoraviruses and their amoebal hosts despite previous literature placing Slo1 channels exclusively in higher animals (Tao and MacKinnon, 2019). From BLAST and jackhmmer iterative database searches no homologous proteins to the alpha subunit of 6v22 were identified for either viruses or amoeba. To address the possibility that sequence divergence had hampered the identification of homologues, a structural search was done using Foldseek to search the alpha subunit of 6v22 against the AlphaFold Database (Varadi *et al*., 2022), restricting the search to viruses and amoebozoa. The top hit in the AFDB-proteome database with an e-value of 6.04e-50 was a model of UniProt entry Q55CU6 from slime mold *Dictyostelium discoideum*, a soil-dwelling amoeba. In the AFDB50 database, containing a 50% sequence identity, redundancy-reduced version of the entire AFDB resource, further strong hits matching both transmembrane and soluble domains of the alpha subunit, were found in other amoebae including *Dictyostelium purpureum* (best match UniProt code F0ZR89, e-value 2.11e-45) *Tieghemostelium lacteum* (A0A151Z2J9, 1.94e-43), *Heterostelium album* PN500 (D3B9R9, 8.28e-43), *Planoprotostelium fungivorum* (A0A2P6MZ64, 1.39e-42) and *Cavenderia fasciculata* SH3 (F4PHS0, 7.19e-41).

Finally, from BLAST database searches *Dictyostelium purpureum* F0ZR89 identified two homologous acanthamoeba proteins, *Acanthamoeba castellanii* str. Neff (best match UniProt code L8HDA6, e-value 1e-35) and *Acanthamoeba castellanii* str. Neff (L8GN72, 1e-05). HHpred analysis of each yielded 100% probability hits against chain C of the PDB entry 6v22 i.e. the alpha subunit of the K^+^ channel. These two amoebal alpha subunits represent the most likely binding partners of the Pandoravirus cluster 2 proteins identified here as presumed beta subunits. The functional consequences of that interaction remain to be determined but membrane depolarisation, as seen for Chlorella virus, is one possibility.

### Cluster 5 encodes a membrane transporter

Cluster 5 contained a total of 26 sequences with lengths ranging from 499 to 618 residues. The representative sequence chosen for Cluster 5 was A0A2U7UDI8 which has 577 residues. The entry for this sequence in UniProt listed 12 transmembrane helices as features of this sequence, a number confirmed with DeepTMHMM (Table 1). The entry also showed that the 12 transmembrane helices were split into two distinct domains of six helices each, separated by a cytoplasmic domain. DeepLoc predicted lysosome/vacuole localisation as most likely, followed by cell membrane localisation. In addition to the expected pandoraviruses, iterative searches with the PSI-BLAST database searches revealed homologues in bacteria: the first of these was the sequence with UniProt code A0A2E7ZQ83 from Myxococcales bacterium with an e-value of 0.003. Many of these BLAST hits match Domain of Unknown Function (DUF5690) which also aligns to the cluster 5 representative sequence. Interestingly, the Pfam website suggested that pandoraviruses are the only viruses to contain DUF5690, which is most commonly found in bacteria. The Pfam database annotates DUF5690 as a member of the Clan MFS. The major facilitator superfamily (MFS) is one of the two largest families of membrane transporters found on Earth (Pao, Paulsen and Saier, 1998) MFS members, like A0A2U7UDI8, typically have 12 transmembrane helices but some have 14 (Drew *et al*., 2021)). HHpred was used to search for homologues of known structure in the PDB. HHPred produced hits to the PDB spanning across both the N and C-terminus, with strongest hits across the whole molecule to (MFS)-fold transport proteins. The strongest HHPred hit was to the MFS Melibiose carrier protein (PDB: 7l17-A, Prob = 98.49%). Full-length matches were additionally seen to MFS members such as rat vesicular glutamate transporter 2 (PDB: 6v4d-A) and glycerol-3-phosphate transporter from *E. coli* (PDB: 1pw4-A).

AlphaFold2 was used to generate a protein model of the representative sequence. The model was processed by removing regions of low quality prediction, likely intrinsically disordered regions present at the N-and C-termini (residues 1-54 and 508-577, respectively). The model displayed the expected 12 transmembrane helices and featured a connecting region in the form of an unstructured loop/α-helix, which separated the two domains (Fig 4). This connecting region (linker) is a common feature of MFS transport proteins. DALI found many strong hits to the model in the PDB, with Z scores reaching a maximum of 29.1 (Table 2). All strong hits generated were to transport proteins, particularly proteins belonging to the MFS, including the *E. coli* D-galactonate:proton symporter (6e9o), Z score: 29.1. D-galactonate transporters belong to the MFS fold (Leano *et al*., 2019)

**Figure 4.**
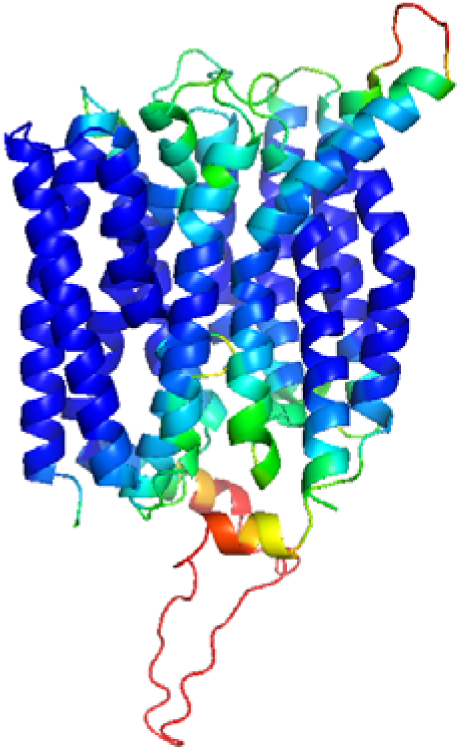
AF2 model in the closed (occluded) conformation: low confidence, likely intrinsically disordered termini are not shown. Standard AF2 colouring was applied. Blue regions represent areas of high confidence, red regions represent areas of low confidence.

Like many families of transporters, MFS proteins are known to have multiple biologically significant conformations. In the case of the MFS these are outward-open, outward-occluded with bound substrate, occluded with substrate, inward-occluded with substrate, inward-open and occluded with no substrate (Drew *et al*., 2021). AF2 can sometimes be induced to sample such alternative conformations by reducing the depth of the multiple sequence alignment used as input (512:1024), but attempts to model other conformations by that route here were unsuccessful.

Despite their rarity, transport proteins encoded by viruses are known to be functional. This includes chloroviruses, which are large dsDNA viruses like Pandoraviruses. Chloroviruses feature known ion channel transport proteins, including potassium and calcium ion channels, which have recognised functional roles during early viral infection (Thiel *et al*., 2015). Another example is the *Ostreococcus tauri* virus RT-2011 that infects the marine alga that has been shown to encode a transporter protein that mediates nutrient uptake from the environment (Monier *et al*., 2017). In the current literature, pandoraviruses are not reported to encode such transporters, but these precedents lent strong credibility to our annotation of cluster 5 as a transporter for an as-yet unidentified substrate.

In an attempt to characterise potential substrates further, electrostatic analysis was carried out in PyMOL using APBS (Fig 5). This revealed a mainly internal positive charge at the interface between the N- and C-terminal domains, the presumed site of passage of a transported substrate, possibly indicating that the substrate for this specific transporter is negatively charged.

**Figure 5.**
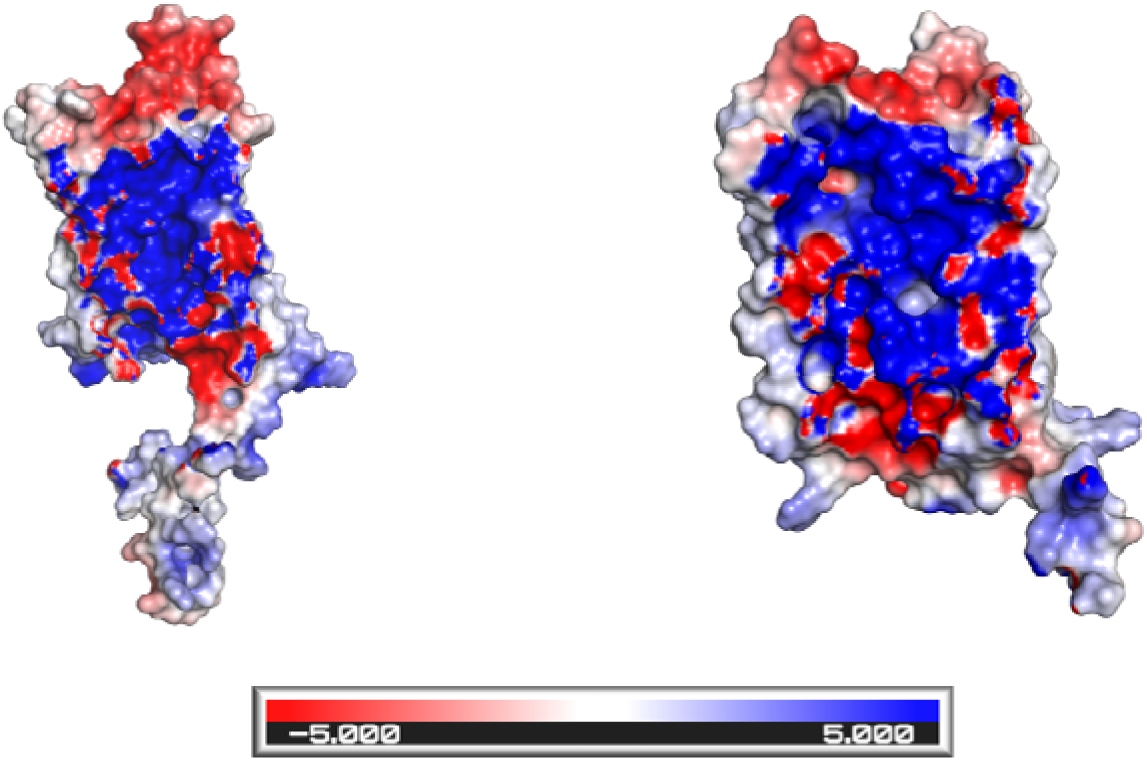
APBS electrostatic analysis of the domain interface of the cluster 5 representative A0A2U7UDI8 showing in open-book representation a) the N-terminal domain and b) the C-terminal DUF5690 domain. Blue indicates positive charge, red negative charge.

Finally, phylogenetic analysis of cluster 5 revealed three confident clades, each containing one sequence from each of the four Pandoraviruses considered here (Supp Fig 1). Inspection of the alignment revealed 20 amino acids in the reference model to be of particular interest since they were conserved within each clade but different between the three clades (Supp Table 1). The positions of these 20 amino acids were mapped on the model to suggest possible functional implications. Interestingly, two amino acids, L79 and A334 were sited at the central binding cavity (Fig 6) suggesting that the characteristic, clade-specific substitutions found in other members of cluster 5 - L79 replaced by F or Y; A334 replaced by G or V - could alter the substrate specificity of the transporter.

**Figure 6.**
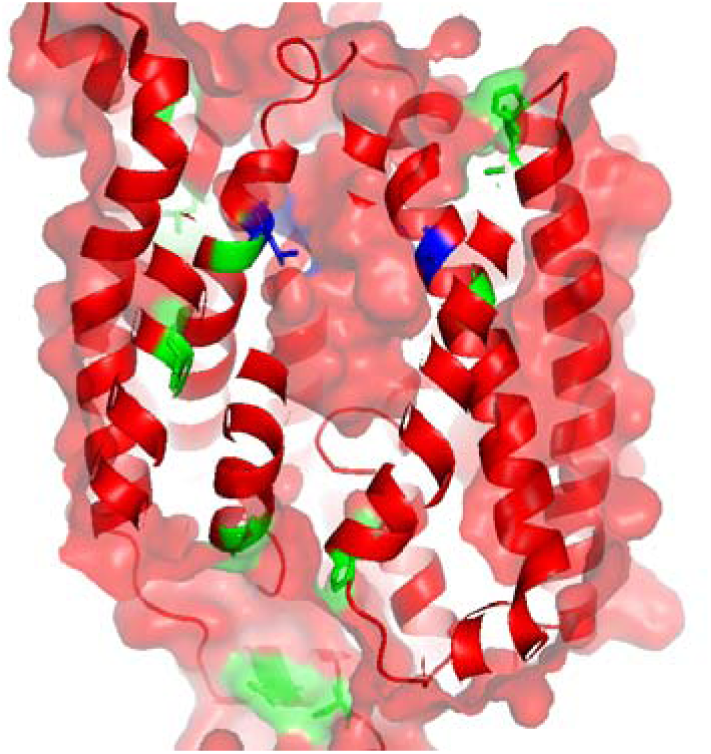
The 20 clade-specific positions identified by the phylogenetic analysis are mapped to the AF2 model of the protein shown in surface representation. Blue positions indicate the two amino acids L79, left, and A334, right that line the central binding cavity. Green indicates the other 18 positions.

### Functionally cryptic and structurally novel proteins

In many cases, AF2 produced models of high predicted quality but matches to the familiar folds, particularly helical repeats, that resulted spanned a wide array of potential functions and little to nothing could be said about the function of the pandoravirus proteins. In other cases, high quality models lacked close structural neighbours in the PDB or databases of models: in many of these cases too, only minimal functional annotation could be carried out

#### Cluster 1

There are 58 proteins contained within this, the largest cluster, from which A0A2U7UAA1 with 289 residues was chosen as the representative protein. Further analysis revealed this predicted cytoplasmic protein, with no major regions of predicted intrinsic disorder, to have no apparent homologues outside Pandoraviruses (Table 1). The AF2 model (Fig 7) was largely highly confident, but for the N-terminal region, and revealed a single globular domain composed largely of a-helices but with a four-strand β-sheet.

**Figure 7.**
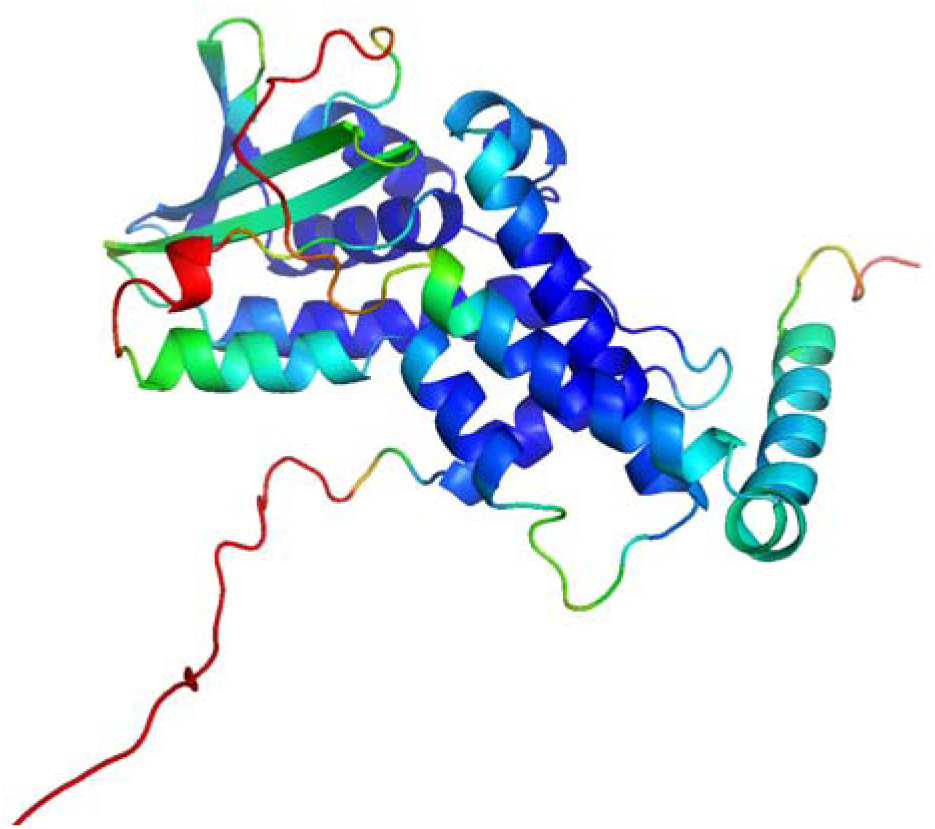
AF2 model of the protein with UniProt code A0A2U7UAA1, the chosen representative protein for cluster 1. Colouring is according to the AF2 pLDDT confidence values, with blue representing high confidence and red low.

DALI hits in the PDB tended to match either the beta-sheet region or the bundle of α-helices but not both. None of the matches scored particularly strongly (Table 2): the highest scoring match, with a z-score of 6.8, involved the β-sheet region, and was for a match to an uncharacterised protein YkuJ (PDB code 2ffg, unpublished). Another hit matched the bundle of α-helices to the RAS exchange (REM) domain of Ras-specific exchange factor RasGRP1 (Chain 4l9m-A) and had a z-score of 4.1 (Fig 8a). Although a weak match, this was deemed worthy of further attention since interference in host signalling pathways is such a common feature of virus-host interaction (Alto and Orth, 2012).

**Figure 8.**
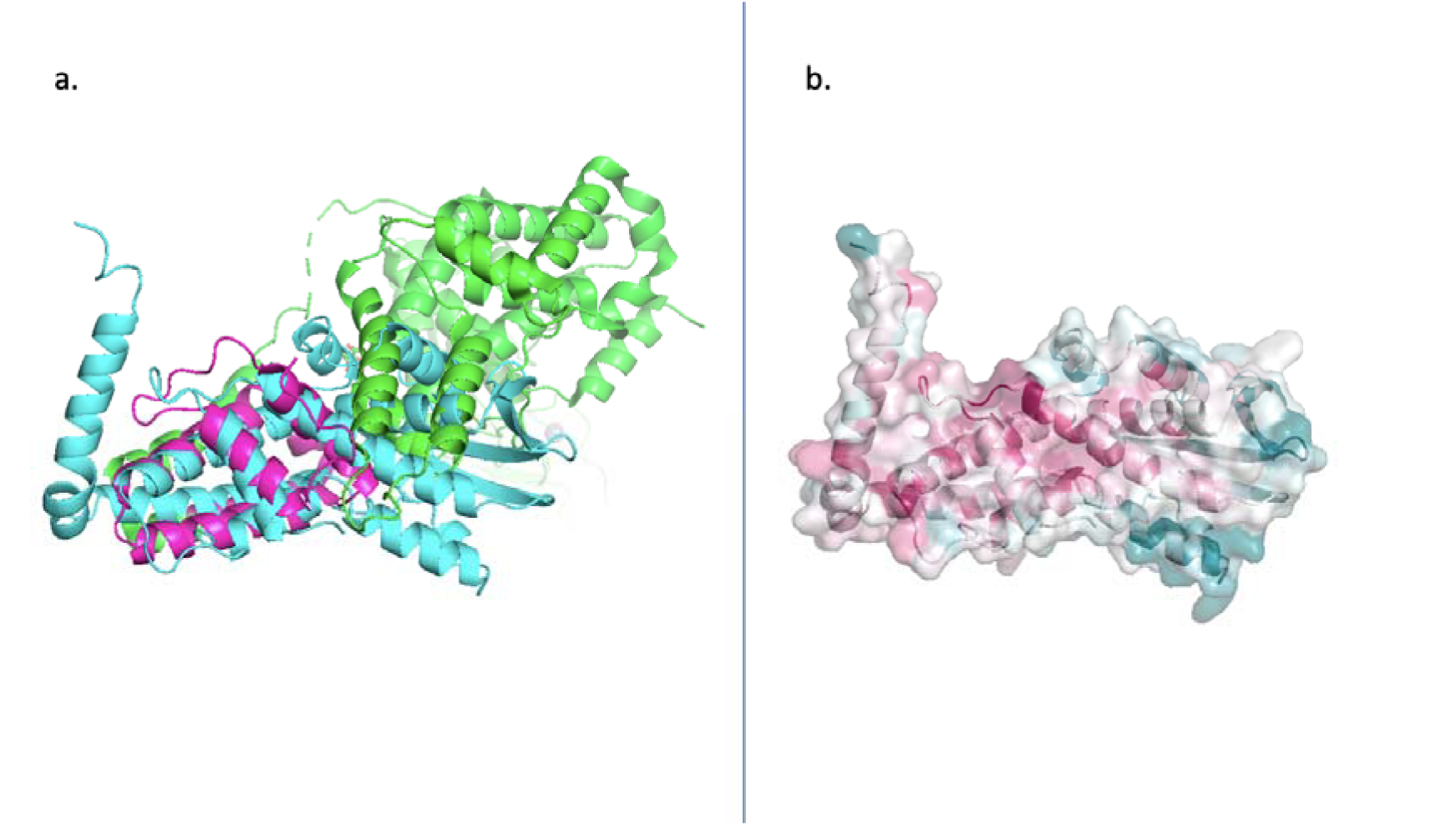
The image on the left (a) is the DALI alignment of A0A2U7UAA1 model (cyan) and RasGRP1 (green) with the REM region highlighted in magenta. On the right (b) is the consurf model of A0A2U7UAA1 with a semi-transparent surface. In each panel low-confidence residues 1-22 are not shown.

Consistent with such a functional role, Consurf indicated high conservation of the area of the AF2 model matching the REM domain (Fig 8b). However, the REM domain activates Ras via binding at a distal binding site (Margarit *et al*., 2003). Examination of the A0A2U7UAA1 model showed that this site was blocked by an α-helix positioned in a way that would prevent Ras binding. Since the helix was not integral to the fold, further modelling was carried out to see if alternative structures positioned the helix differently, potentially liberating a site with a REM-like function. Modelling was done with RoseTTAFold and by giving AF2 shallower MSAs (this can induce the modelling of alternative conformations; ref). The RoseTTAFold model was extremely low confidence and no longer had structural resemblance to the REM domain of RAS-GRP1 while the further AF2 runs also resulted in active site occlusion. Finally, the model was run with the distal Ras and REM domain structures from PDB entry 1NVW as a dimeric template in an attempt to force positioning of the α-helix away from the active site. This also proved unsuccessful, with occlusion still taking place. Thus, while a REM-like function cannot be ruled out, the evidence remains weak and this large cluster remains functionally uncharacterised

#### Cluster 3

Cluster 865 comprised 21 sequences with lengths ranging from 276 to 388 residues. From which the chosen representative was A0A2U7UDC4 with 296 residues. Subcellular localisation prediction using DeepLoc predicted cytoplasmic and nuclear localisations as most probable: other sequence analyses were negative (Table 1). BLASTp searches and iterative searches using jackhmmer revealed that homologues were confined to pandoraviruses. Searches using HHpred produced the strongest hits ankyrin-repeat and subunits and domains, such as Serine/threonine-protein phosphatase 6 regulatory ankyrin repeat subunit B (E-value: 7.3e-14) and Ank_2 with an E-value of 0.00022. AF2 was used to generate a protein model of the representative sequence. N- and C-termini regions of intrinsic disorder (residues 1-50 and 560-596, respectively) were first removed before AF2 structure prediction was run.

Searches against the PDB with this AF2 structure using DALI returned 296 hits with a Z-score greater than or above 7.0 reflecting the frequency with which such α-helical proteins are found in nature. In agreement with the HHpred results, DALI presented hits to a number of ankyrin repeat domains, such as to the human ankyrin repeat domain of Bcl-3: a unique member of the IkappaB protein family. (PDB: 1k1b-A, Z-score: 8.3). Ankyrin repeats are found in many functionally diverse proteins (Mosavi *et al*., 2004; Monier *et al*., 2017)) but exclusively function to mediate protein-protein interactions, suggesting this function for Cluster 3.

FoldSeek was additionally used to search for structural neighbours in the PDB. FoldSeek returned many hits, particularly to Pentatricopeptide repeat (PPR)-containing proteins. The PPR proteins are a large family of eukaryotic RNA binding proteins. Highest PPR-containing protein match was to a mitochondrial Zea mays protein (Prob = 0.80, TM score = 0.40688, UniProt = A0A096RDL4). An RNA binding role may be considered unlikely, however, since the AF2 model of the representative displays strongly electronegative character (Fig 9) while RNA-binding proteins typically possess positively charged surfaces (Shazman and Mandel-Gutfreund, 2008). In particular, viral RNA-binding proteins are commonly positively charged, with less than 20% exhibiting a negative or net zero charge (Ahmad and Sarai, 2011).

**Figure 9.**
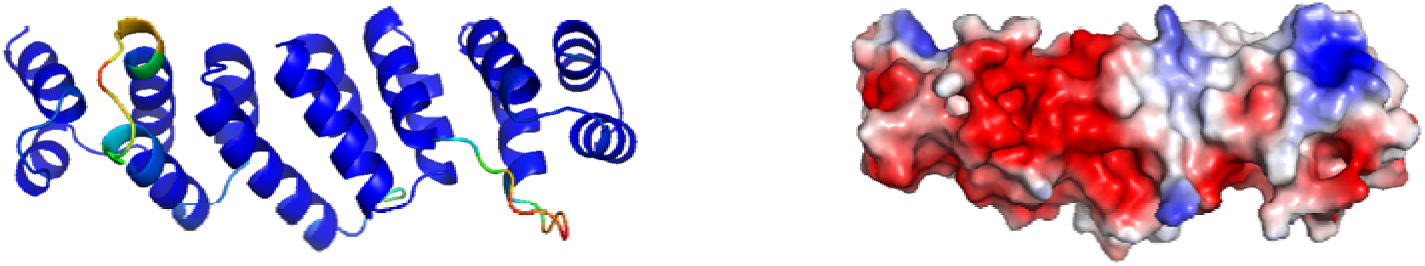
AlphaFold2 model. Left) coloured by pLDDT values, with blue representing high confidence and red low confidence. Consistently low confidence, likely intrinsically disordered termini are not shown. Right) Electrostatic analysis of the AF2 model of the Cluster 3 representative sequence, coloured red (negatively charged) to blue (positively charged)

Furthermore, modular interactions of PPR proteins with RNAs occur through specific amino acid recognition codes within PPR protein motifs (Yagi *et al*., 2013) Such amino acid recognition codes were not identified in Cluster 3. Additionally, Foldseek likewise returned many hits to Ankyrin Repeat region-Domain containing proteins, the highest ankyrin repeat match being against UniProt: I1LNB6, (TM score = 0.3717, prob = 0.82). The overall best FoldSeek match against the AFDB was an unplaced genomic scaffold, Prob = 0.99, TM score = 0.44678, A0A0D2DAR4.

The electrostatic analysis suggested a function more like that of ankyrin repeat domains i.e. protein-protein interaction than an RNA-binding function. Ankyrin repeats are particularly rare amongst viruses (Odon *et al*., 2018) but have been documented in poxviruses, which may have acquired them through horizontal gene transfer (HGT) (Odon *et al*., 2018). Both poxviruses and pandoraviruses are megaviruses within the Nucleocytoplasmic Large DNA Viruses (NCLDV) family (Krupovic, Yutin and Koonin, 2020). It is therefore plausible that pandoraviruses may have acquired ankyrin repeats through the same mechanism. While the functions of most poxviral ankyrin proteins remain to be elucidated, viral ankyrin proteins such as ANK/BC proteins have been discovered and function as immunomodulatory proteins that inhibit innate host immune responses (Odon *et al*., 2018). Such a function is unlikely given the unicellular amoebae hosts of pandoraviruses, meaning that the identity of any interaction partner must await experimental study.

#### Cluster 4

This cluster contained 27 protein sequences around 250 residues in length, except for A0A291AU05 which had around 100 extra residues at the N-terminus. The representative A0A2U7U8Z7 (*P. quercus*) was of length 228 residues, contained a signal peptide (residues 1-31), and was predicted to be secreted by DeepLoc (Table 1). HHpred found no significant matches in the PDB or Pfam. Five iterations of PSI-BLAST found 57 hits in Pandoraviruses. A wide variety of other megaviruses also contained clear homologues, including Mimiviruses, Powai lake megavirus, and Bandra megavirus. None of these hits were functionally annotated.

AF2 produced a highly confident single domain globular structure rich in β-sheets (Fig 10). DALI found many strong hits matching beta-trefoil structures, the highest Z-score being 15.7 (Supp Table 2). The top 100 hits were investigated, of which DALI sequence alignments showed conserved β-sheets and loop structures, and a total of seven Pfam domains were present. Of these domains, the highest Z-scoring entry for each was taken as a representative for further analysis (Supp Table 2) with the aim of determining whether the Cluster 4 function might be related to those of any of the DALI hits. FoldSeek was also used to search the AFDB (data not shown) yielding similar hits to DALI, but no major distinct functional hypotheses.

**Figure 10.**
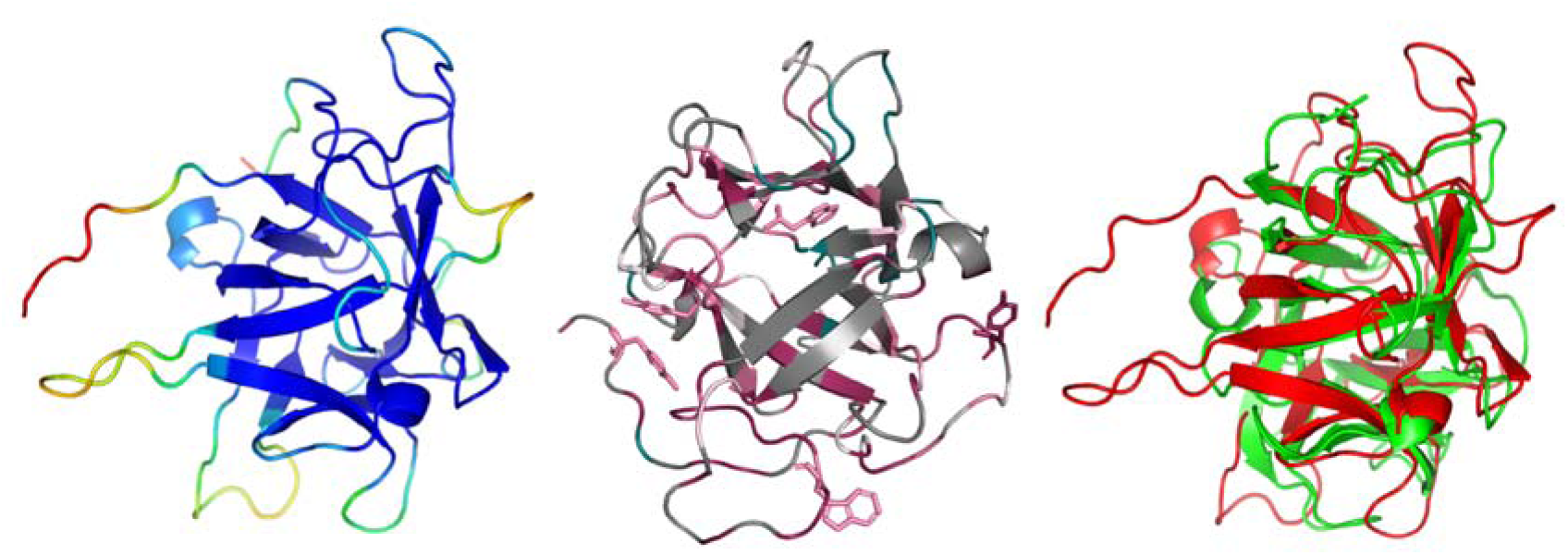
Left: AF2 model of A0A2U7U8Z7, coloured by confidence. Centre: Consurf mapping of conservation on A0A2U7U8Z7 (coloured as Consurf, with 94 residues having insufficient information coloured in grey), with highly conserved aromatic Trp and Tyr residues shown. Note the particularly well conserved loop from residues 167-178 in the lower part of the panel, and the potential for outward facing aromatic residues for carbohydrate binding. Right: Superposition of A0A2U7U8Z7 (red) and 3mal-B (green).

The functional contexts, or an absence of specific characteristics immediately argued against a functional connection to some DALI matches. For example, 3im5-B is domain B of the N-terminal region of the Ryanodine receptor 2, and 3v0b-B is the nHc region of the large multi-domain NTNHA protein, both of which have structural roles involved in interdomain interactions within the receptor, and also protein-protein interactions with specialised *Clostridium botulinum* neurotoxin in the case of 3v0b-B (Lobo and Van Petegem, 2009; Gu *et al*., 2012; Kimlicka *et al*., 2013). As a single domain protein, the cluster 4 representative cannot have this function. Similarly, 5hpz-B (PF00197: Trypsin and protease inhibitor) binds chlorophyll, yet cluster 4 contains none of the required conserved Pro residues and *Acanthamoeba spp.* are not photosynthetic (Bednarczyk *et al*., 2016). Additionally, the wider protease inhibitor functionality of PF00197 are difficult to assign to Cluster 4, as usually conserved Arg, Asn and Lys residues, alongside two conserved disulfide bridges that support and form the active site of an inhibitory loop, are absent (Bednarczyk *et al*., 2016; Bendre, Ramasamy and Suresh, 2018).

A potential carbohydrate binding function was further investigated, as four of these domains are associated with carbohydrate binding, represented by: 3win-B (PF14200: Ricin-type beta-trefoil lectin domain-like), 3uj4-A (PF02815: MIR domain), 1jly-B (PF07468: Agglutinin domain), and 3n7j-A (PF07951: Clostridium neurotoxin, C-terminal receptor binding). 3win-B was discounted as A0A2U78Z7 matches the non-carbohydrate binding domain of 3win-B, thought to be involved in interactions within the specialised Haemagglutinin complex of *C. botulinum* neurotoxin (Nakamura *et al*., 2008, 2011; Amatsu *et al*., 2013), and also lacks the conserved QxW Ricin B Lectin motif (Rotskaya *et al*., 2021). The remaining three: 3uj4-A, 1jly-B and 3n7j-A bind inositol 1,4,5-trisphosphate, T-antigen disaccharide, and both sialic acid and a ganglioside respectively. However, most key conserved binding residues were absent from the Cluster 4 alignment save for Tyr50, which was still not universally conserved (Yoshikawa *et al*., 1996; Transue *et al*., 1997; Karalewitz *et al*., 2010; Strotmeier *et al*., 2010; Seo *et al*., 2012; Lee *et al*., 2016).

More broadly, beta-trefoils can potentially bind a diversity of carbohydrates due to their low shared sequence similarity (Žurga *et al*., 2015; Acebrón *et al*., 2023). Whilst binding typically takes place via loop structures, which loops may be involved differs between trefoils, even when binding similar ligands (Transue *et al*., 1997; Acebrón *et al*., 2023). Trp and Tyr are known to be generally important in carbohydrate binding sites (Armenta *et al*., 2017), and Consurf identified five highly conserved residues: Tyr2, Tyr50, Trp60, Tyr95 and Trp171, of which Trp171 falls within a well-conserved loop (Fig 10). However, in the cluster 4 alignment, only Tyr50 is somewhat conserved so a carbohydrate binding functionality lacks strong support for Cluster 4.

The highest scoring single domain protein in the DALI results was 3mal-B which matches Pfam entry PF02815 MIR (Fig 10). 3mal-B is Stromal Cell-Derived Factor 2-Like 1 protein (SCDF2L1) from *Arabidopsis thaliana* (UniProt: Q93ZE8), which has an ER retention motif and a signal peptide. SCDF2L1, along with its homologue Stromal cell-derived factor 2 (SCDF2), are highly conserved amongst animals and plants, and are upregulated in the eukaryotic unfolded protein response (UPR) in the ER. SCDF2L1 forms a complex with the ERdj3 chaperone, promoting the chaperone’s activity and reducing protein aggregation (Schott *et al*., 2010; Fujimori *et al*., 2017; Hanafusa, Wada and Hosokawa, 2019).

Interestingly, while no ER retention signal was predicted for A0A2U7U8Z7, SDF2 in mice also lacks an ER retention signal, being instead held in the ER by interaction with other UPR proteins (Lorenzon-Ojea *et al*., 2014). Since viral proteins can initiate the UPR to facilitate intense viral protein production e.g. (Ye *et al*., 2011), it seemed plausible for cluster 4 to have a related role. Thus, to investigate a possible SCDF2L1 functionality for cluster 4, the A0A2U7U8Z7 structure prediction, 3mal-B and an AF2 model of mouse SCDF2L1 (*Mus musculus;* Q9ESP1, with signal peptide residues 1-28 removed) were analysed by CSM-potential. Each structure had regions strongly predicted as protein-protein interaction interfaces but the Pandoravirus structure was positioned differently in a way that could not be accounted for by the threefold pseudo-symmetry of beta-trefoil structures (Kirioka, Aumpuchin and Kikuchi, 2017; Blaber, 2020) (Supp Fig 2).

To explore a potential SCDF2L1-related function further, a local AF2 installation was used to explicitly model the interaction of the cluster 4 model and the same SCDF2L1 structures with corresponding ERdj3 chaperones. The chaperone sequences in mouse (UniProt: Q99KV1) and Arabidopsis (UniProt: Q9LZK5) were identified by text searches in UniProt. HHpred could find no ERdj3 homologues in *P. quercus*, but reciprocal BLAST with *A. thaliana* ERdj3 (UniProt: Q9LZK5) identified a putative ERdj3 orthologue (UniProt code: L8H8X3) in *A. castellanii* Neff strain sharing around 41% sequence identity. While convincing heterodimers were built of the mouse and Arabidopsis pairs, the model of cluster 4 in complex with the amoebal chaperone was lower confidence and unconvincing (Supp Fig 3). Overall, while roles in the UPR, and/or in carbohydrate binding/recognition might be possible, Cluster 4 remains functionally uncharacterised.

#### Cluster 6

The largest cluster containing structurally highly novel domains was cluster 6 which comprised 25 sequences with around 700 residues. Twelve of the sequences matched DUF5768 (PF19072) in their Pfam entries; the others contained no Pfam domain annotations. Sequence-based predictions (Table 1) were largely uninformative, but suggested the protein to be cytoplasmic. Iterative database searches with PSI-BLAST and JackHMMER showed that clear homologues were confined to pandoraviruses and again revealed DUF5768, for which there is no information in the literature, as the only matching entry in Pfam or InterPro.

AF2 produced a largely confident model prediction with three distinct domains, each consisting of several β-sheets surrounded by α-helices (Fig 11). These domains visually comprised residues 1-263 (domain A), 264-469 (B), 470-691 (C), an assignment largely consistent with automated SWORD2 annotations (domain A: 1-265 plus 363-377; domain B: 266-362 plus 411-468; domain C: 469-691 plus 378-410). The domains are connected by several large loop structures which have lower confidence. The superficial similarities between the domains prompted us to consider potential distant homology but DALI all-against-all Z-scores of 2.3-2.4 disprove this, as did differences in topology revealed by closer examination.

**Figure 11.**
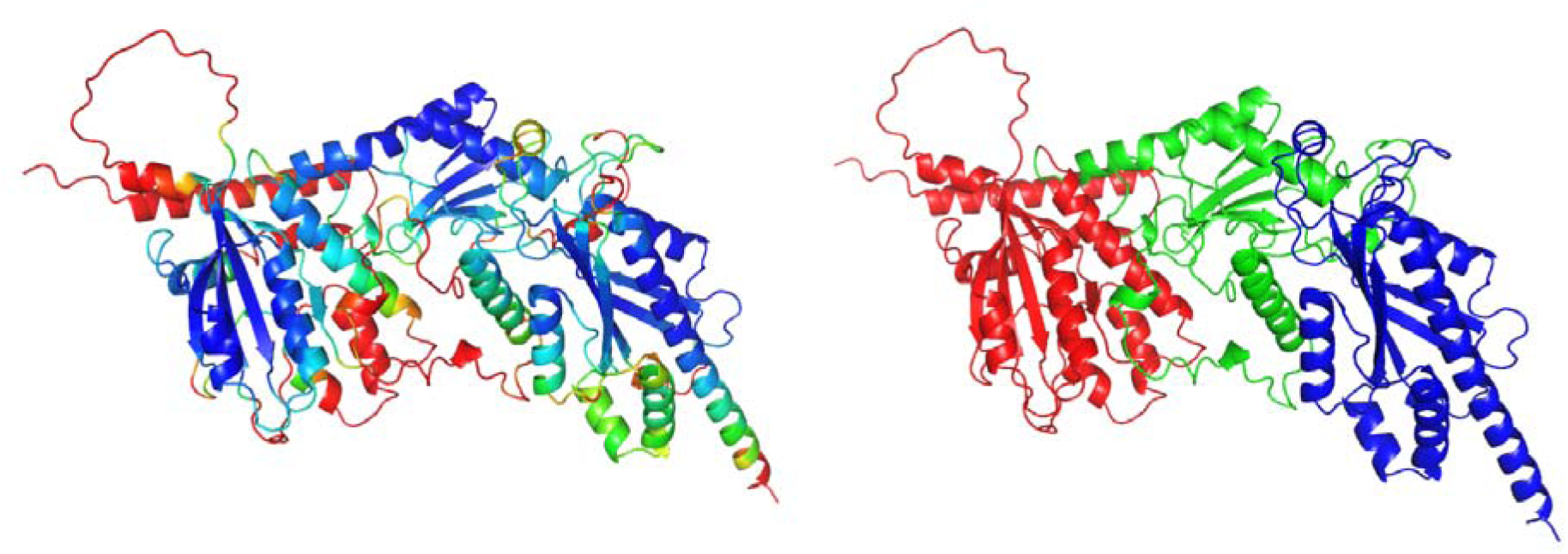
Left: AF2 model of A0A2U7UCR8 coloured by confidence. Right: The three domains of A0A2U7UCR8: Domain A (red), Domain B (green) and Domain C (blue).

Table 2 shows the structural neighbours in the PDB (wwPDB consortium, 2019), AFDB (Varadi *et al*., 2022) and ESM Metagenomic Atlas (Lin *et al*., 2023) for the three domains. Domain A gave no PDB matches with a Z score higher than 5.7 and was not investigated further. The strongest DALI Z score was 7.8 for domain B against 1s28 chain A, the structure of a type 3 secretory chaperone (T3SC) (Fig 12). The cluster 6 domain B model differs significantly from that topology yet, given the intense protein production that occurs during pandoravirus infection, a protein chaperone function initially seemed plausible. However, detailed examination suggested it lacked key functional features. For example, the surface of 1s28-A is significantly negatively charged, and contains two hydrophobic patches that interact with the effector (Singer *et al*., 2004). However, domain B of the model is not negatively charged, and no corresponding hydrophobic patches are present. In addition, the alpha-2 helix of the T3SC contributes to dimerization (Singer *et al*., 2004), but when 1s28-A is aligned with domain B, this helix becomes buried within domain C, and so the same manner of dimerization - considered essential for function (Parsot, Hamiaux and Page, 2003) - is implausible for domain B. DALI found ten other PDB hits to T3SCs or T3SC-like folds for domain B, with Z-scores 5.0-5.6, eight of which function as T3SCs, and all having a characteristic topology (Norais *et al*., 2013). Finally, CSM-potential does not strongly predict protein-protein interaction sites for domain B. Taken together, these results suggest that domain B does not have the chaperone function and remains functionally cryptic.

**Figure 12.**
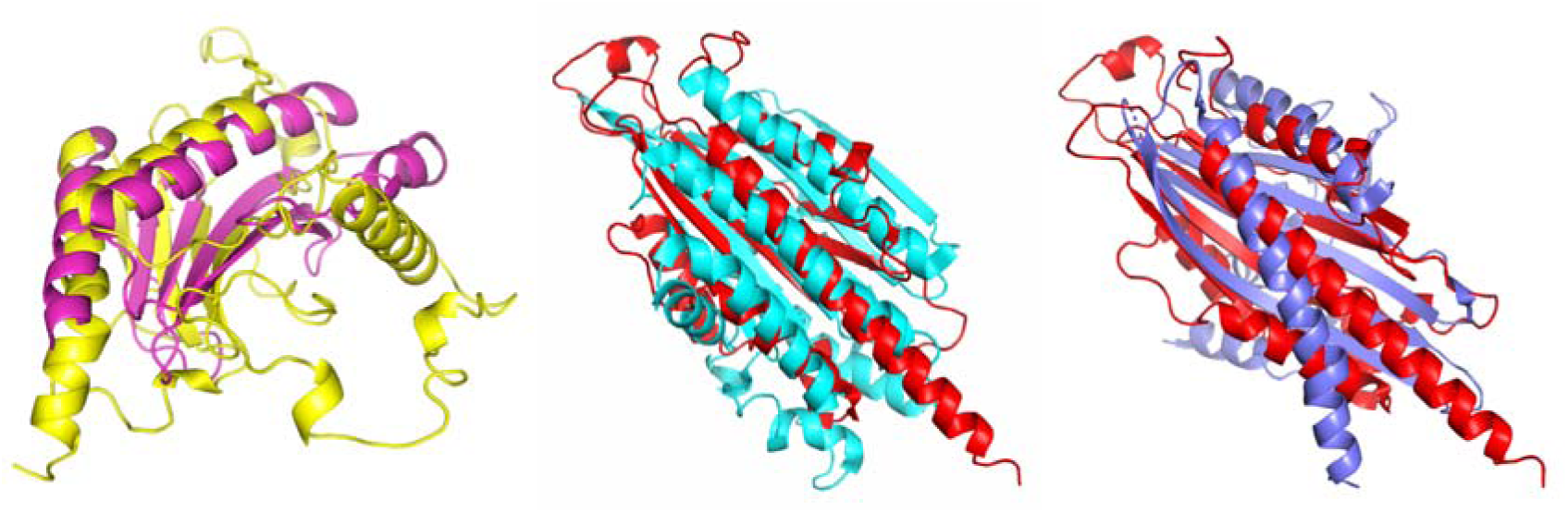
Left: superposition of Domain B (yellow) and 1s28-A (magenta). Centre: superposition of Domain C (red) and 3dwr-A (blue), the highest scoring DALI hit (Z=7.0). Right: superposition of Domain C (red) and 3lk4-B (lavender), the highest scoring actin-capping protein DALI hit (Z=6.7).

The three highest structural matches for Domain C were oxidoreductase enzymes: two coproporphyrinogen III oxidases (3dwr-A, Z=7.0, Fig 12; 1txn-B, Z=7.0) and a phycoerythrobilin synthase (2vck-C; Z=6.7). However, conserved Histidine, Arginine, Aspartic acid and Proline residues crucial for the activities of these enzymes could not be identified in Domain C; indeed, none of these residues are conserved across the entire MAFFT cluster 6 alignment, likely discounting this type of function (Dammeyer, Hofmann and Frankenberg-Dinkel, 2008; Silva and Ramos, 2011).

Since divergent actin homologues are present in megaviruses (Da Cunha *et al*., 2022), the weak structural matches of Domain C to actin capping proteins of Z scores: 6.6-6.7 were intriguing (Fig 12). However, closer inspection again demonstrated the absence of key features. In the actin capping heterodimer, an extended amphipathic helix at the C-terminus of both subunits is important for interacting with actin, and is strongly conserved amongst all capping proteins (Hernandez-Valladares *et al*., 2010); (Funk *et al*., 2021; Bendes, Kursula and Kursula, 2022). Whilst sequence conservation in this “tentacle” is low across eukaryotic capping proteins, there is a generally conserved hydrophobic pattern of leucines (L258, L262 and L266 in *Mus musculus*; (Kim, Cooper and Sept, 2010). Whilst there is a helix that seems to correspond to the tentacle in Domain C, it has low sequence conservation according to Consurf, does not contain the pattern of leucine residues, and is not clearly amphipathic. Additionally, AF2 did not confidently dimerise Domain C, and did not predict a dimer interaction that resembled that of the actin capping dimer. Nor could AF2 build a convincing model of the cluster 6 representative sequence in complex with actin homologues identified in *Acanthamoeba castellanii* (not shown) (we could not detect actin homologues in the Pandoravirus itself). Again, in sum, the actin capping function, though plausible, seemed to lack support.

With uninformative structure matches, functional sites were also sought by using Consurf to map sequence conservation onto the AF2 model. This was done both with a MAFFT alignment of cluster 6 members and using the Consurf server’s inbuilt database search. However, no likely functional sites could be identified using this method or STP leaving the proteins in cluster 6 functionally uncharacterised.

#### Cluster 8

The protein with UniProt code S4VVH3, having 275 residues, was chosen to represent the 22 proteins in this cluster. Further database searches showed apparent homologues only in Pandoraviruses and sequence analysis revealed no apparent features such as intrinsic disorder or signal peptide (Table 1). Unusually, a mitochondrial subcellular localisation was predicted for this protein which gave no significant PDB or Pfam hits by HHpred. The AF2 model (Fig 13) was,on the whole, quite confidently predicted, but with some uncertainty at the N-terminus, as well as within a small section that joins two α-helices.

**Figure 13.**
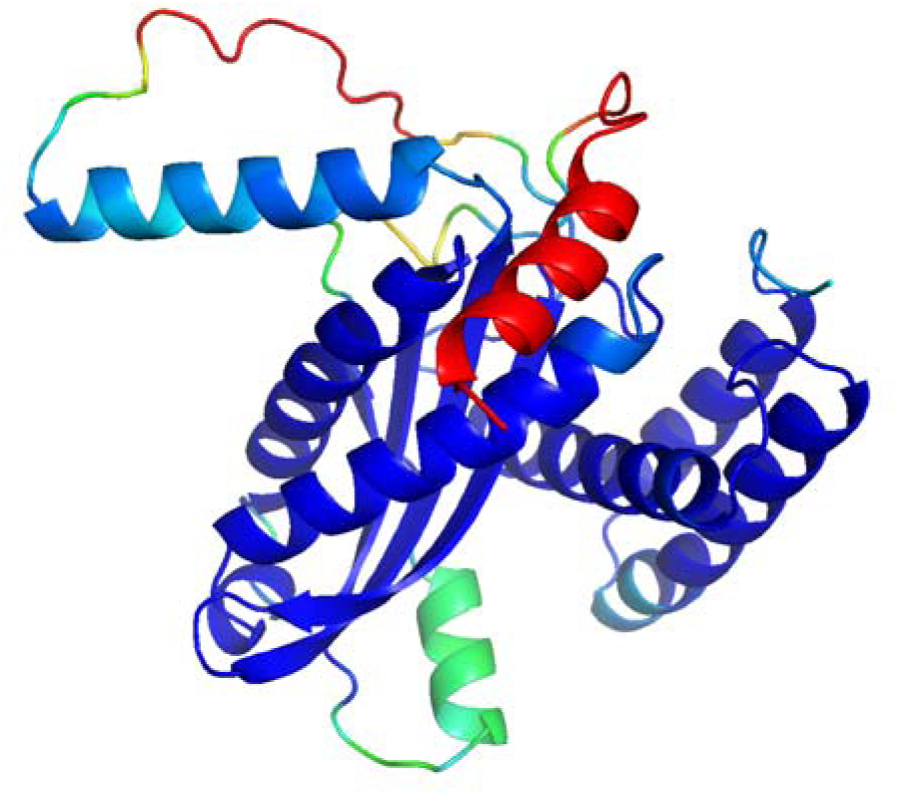
AF2 model of representative protein of cluster 870. Colouring is by AF2 pLDDT confidence values.

The top z-scores returned by a DALI search against the PDB were quite strong, with the top two being 9.1 and 8.9, for stomatin (chain 3BK6-A) and Flotillin 2 ( chain 1win-A), respectively. FoldSeek recovered the same similarities to stomatins and flotillins in the search against the PDB but gave stronger hits to AFDB entries for prohibitins. Prohibitins, stomatins and flotillins all share a conserved domain, namely the Band 7/SPFH/PH*b*-domain containing proteins (Morrow and Parton, 2005; Ibarrola-Vannucci *et al*., 2021). Intriguingly, prohibitins localise to the mitochondrial membrane, matching the Deeploc predicted localization for cluster 8.

The mitochondria could be an important target since it is involved in many important processes such as programmed cell death, cell proliferation, production of ATP, as well as lipid, amino acid and nucleotide synthesis (Ohta and Nishiyama, 2011). Interrupting or hijacking these processes in amoeba is a known mechanism by which viruses aid their propagation and replication (Ohta and Nishiyama, 2011). Although their molecular function remains unknown, PH*b* domains have been implicated in a wide variety of mitochondrial processes (Signorile *et al*., 2019). Recent work (Schweke *et al*., 2023) suggests that diverse proteins with the fold share the ability to form membrane-associated ring structures. It is therefore tempting to propose that cluster 8 proteins could do the same, although it should be noted that they lack the oligomerisation domain, commonly found in these proteins, and also the N-terminal transmembrane helix which assists mitochondrial localisation in other PH*b* domains (Oyang *et al*., 2022). Similar ring structure formation would thus require alternative mechanisms for self-association and membrane binding.

#### Cluster 9

Cluster 9 contained 21 protein sequences between 283 and 416 residues long. Sequence analysis suggested that the cluster representative is a cytosolic protein with no apparent homologues outside pandoraviruses. Furthermore, HHPred showed that it bore no obvious homology to Pfam families or proteins of known structure. Modelling with AF2 produced a structure (Fig 14a) that was highly confident in the C-terminal alpha/beta portion from residues 150-283 but less so in the α-helical remainder from residues 1-149.

**Figure 14.**
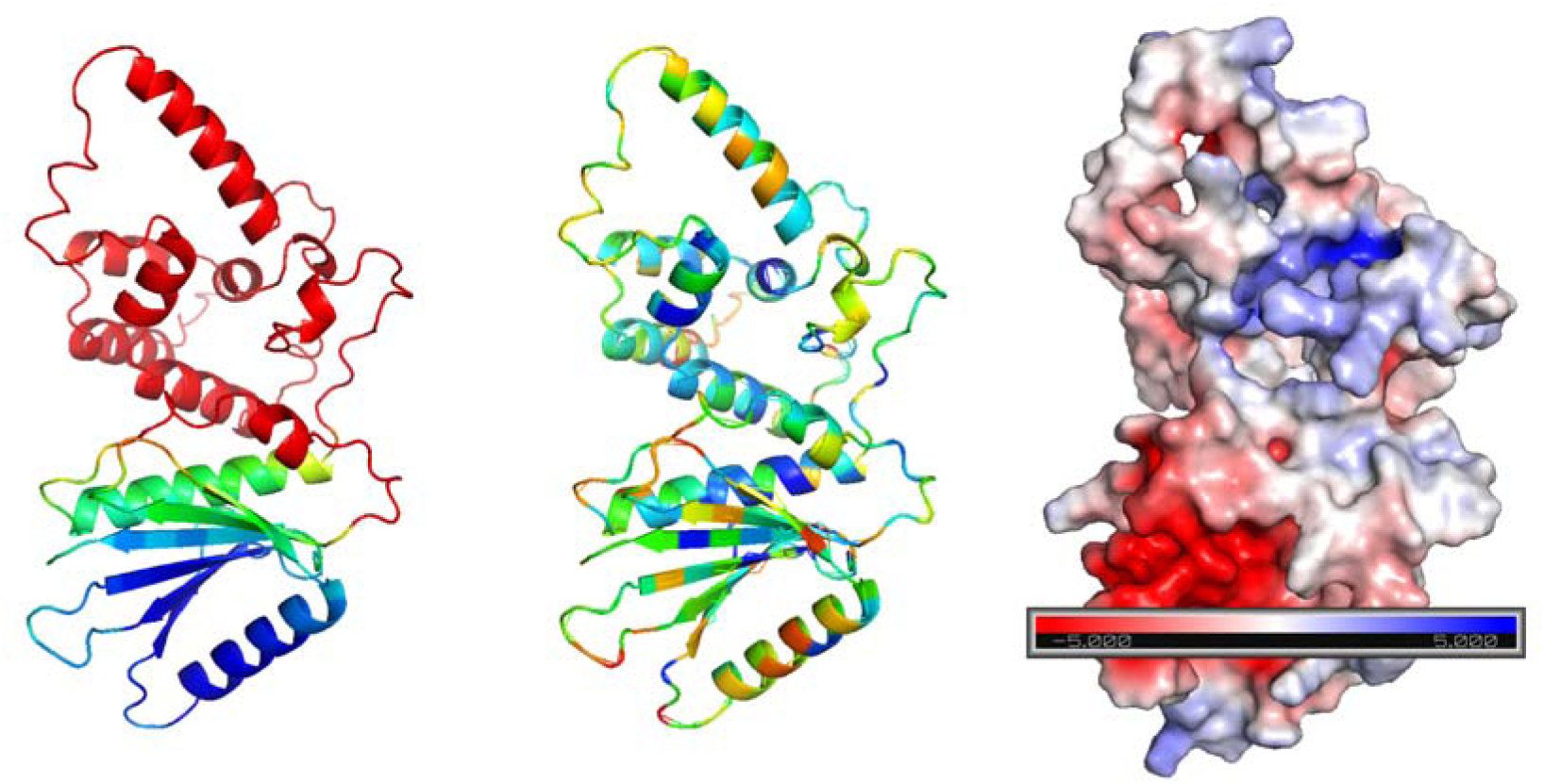
Structural analysis of A0A2U7U902. (a) AF2 prediction, pLDDT used to colour the prediction by confidence with blue being of highest confidence and red of lowest confidence. Highlights the C-terminal domain from residues 150-283 to be of high structure prediction shown in blue. (b) ConSurf conservation analysis where blue indicates conservation and red unconserved residues. (c) APBS electrostatic analysis, red highlighting a negative charge and blue a positive charge on the surface of the protein. The same C-terminal domain displays a distinct area of negative charge in red.

Searches of structure databases, focussing on the more confident domain did not produce strong hits (Table 2) of similarity with at most only half the residues in the domain aligning. The top hit was Pseudopilin EpsI with a Z score of 5.3. The AlphaFold Database gave a top hit of Snoal-Like domain containing proteins with a TM score of 0.48. SnoaL-Like domains are from a family of polyketide cyclases (Sultana *et al*., 2004) this function seems to be unlikely and with a low TM score is indicative only of shared topology, not of strong structural similarity and hence potentially shared function.

With structure matches again functionally uninformative, structure-based model properties were analysed. Electrostatic analysis highlighted a distinct negative cluster within the domain of highest confidence in structure prediction (Fig 14c); however, this is not of particularly high conservation (Fig 14b). Furthermore, pockets located with CastP were relatively small. One had a pronounced negative electrostatic potential but ConSurf analysis did not show it to be highly conserved, nor did STP analysis suggest it had any pronounced ligand binding capability. Overall, the function of proteins in this cluster remains mysterious.

#### Cluster 10

Cluster 10 contained 21 sequences ranging in length between 87 and 320 residues long. S4VZQ1 (234 residues long) was chosen as a representative for this cluster and was predicted to reside in the Endoplasmic Reticulum. Sequence analyses predicted two transmembrane helices between residues 46-68 and 72-94 but no further features (Table 1). Database searches showed that apparent homologues were confined to Pandoraviruses. HHPred was used to search for distant homologies and returned no significant hits.

Alphafold2 was used to model S4VZQ1 (Fig 15). The model displayed two α-helices, corresponding to the transmembrane predictions, and a globular domain containing a β-sheet and shorter helices (residues 124-234). The globular domain showed high confidence scores and was searched against the PDB with DALI. This revealed no significant hits, with the closest match being Mitochondrial Edited mRNA Stability Factor 1 (PDB: 6P5R). (Table 2).

**Figure 15.**
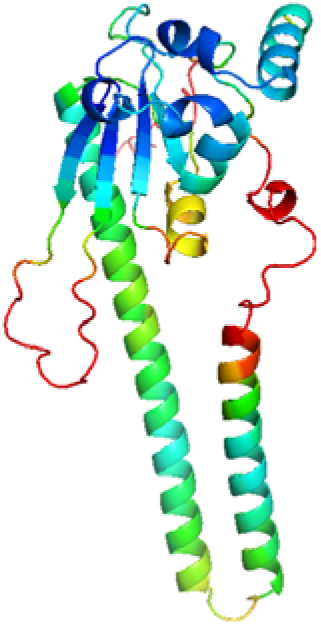
AF2 model of S4VZQ1, coloured by pLDDT from high (ble) to low (red). As shown, S49VZQ1 consists of two transmembrane helices (residues 46-68 and 72-94) and a globular domain (residues 124-234).

Analysis of the globular domain for electrostatics and with STP failed to reveal any strong binding site predictions so the function of Cluster 10 remains cryptic, although its predicted ER location makes it an interesting target for functional characterisation.

#### Cluster 11

Cluster 11 had 16 sequences, ranging between 231 and 626 residues long, from which S4VZY3 was chosen as a representative. S4VZY3 is 441 residues long, has homologues only in other Pandoraviruses, and lacks recognisable features by sequence-based analyses (Table 1). HHPred was used to search for distant homology in the PDB or Pfam but no significant hits were returned. The AF2 model of S4VZY3 (Fig 16) displayed an alpha+beta domain consisting of two β-sheets connected by a short helix (residues 110-208), and α-helical repeat domain (residues 217-441) with an N-terminal region with low confidence scores (residues 1-109). The PAE from AF2 showed that there was very low confidence in the relative orientation of the two domains so, after removal of the low confidence regions, the individual domains were searched against the PDB using DALI.

**Figure 16.**
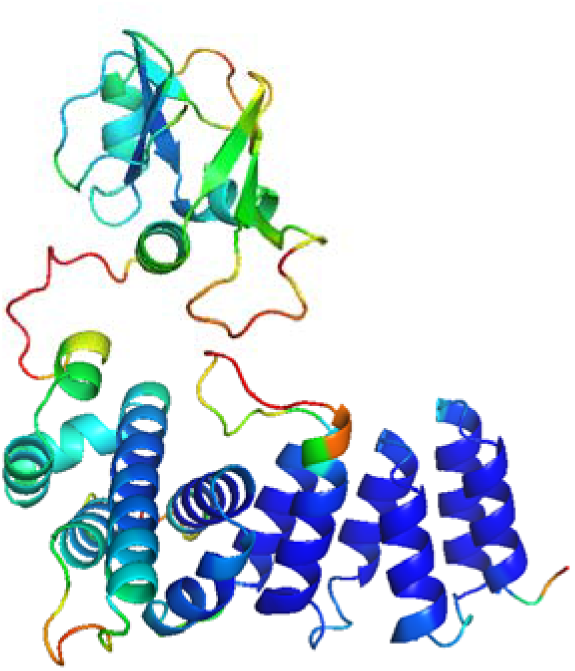
AF2 model of S4VZY3 showing the helical repeat domain (residues 217-441) and αβ domain (residues 110-208). Residues 1-109 are removed due to their low confidence (pLDDT score < 40).

DALI returned several significant hits, mainly due to S4VZY3’s helical domain. No significant hits were returned for the αβ domain. One hit returned was a Pentatricopeptide repeat (PPR) containing protein. PPRs bind to RNA using a positively charged region (Manna, 2015) but the helical domain of S4VZY3 shows strong negative charge and is thus unlikely to bind RNA (Fig 17). Similar to Cluster 3, strong matches against different kinds of α-repeat helix proteins were returned.

**Figure 17.**
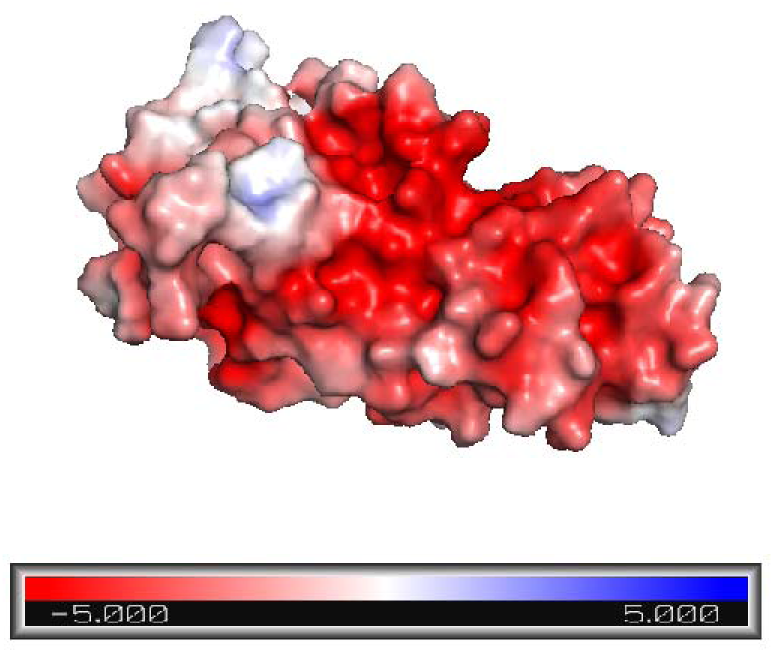
Electrostatic analysis of the α-helical repeat domain (residues 217-441) of cluster 11 representative.

Another significant hit DALI returned was Polyadenylate-binding protein-interacting protein 1 (Paip1, PDB: 6YXJ) Specifically, the helical domain of S4VZY3 aligned with the N-terminal domain of Paip1. The N-terminal domain of Paip1 contains the PAM2 motif (PABP-interacting motif 2) which is known to bind to the C-terminal domain of another protein, PolyA-binding protein (PABP) (Ivanov *et al*., 2016). PAM2 is a 15 residue motif reported to be present in many proteins (Roy *et al*., 2002). However, S4VZY3 did not possess the reported consensus PAM2 sequence, only a similar sequence bearing a large insertion and sited not in the helical repeat region but in a low confidence region.

The quite confidently predicted alpha/beta domain also offered few functional clues, having no close structural neighbours in the PDB. The closest match was the structure of a hypothetical protein PA2021 (Z = 4.2; Table 2). Overall, once again, the function of Cluster 11 remains unclear.

#### Cluster 12

Cluster 12 contained 16 sequences, four from each *Pandoravirus* species, with sequence lengths varying from 166 to 322 residues. S4VS46 (*P. dulcis,* 184 residues), predicted to be located in lysosome/vacuole, was chosen as the representative. In all sequences there were eight conserved cysteine residues numbered 31, 40, 44, 46, 49, 57, 66, and 79 in the S4VS46 sequence. Sequence-based methods predicted a signal peptide at residues 1-18 (removed before modelling), an extracellular region around 19-88, a single transmembrane helix around 89-111, and an intracellular region around 112-184. Disordered regions were predicted between residues 76-80 and 127-175, with an ANCHOR2 predicted disordered binding region at 139-184. Within the ANCHOR2-predicted region, considering other requirements for some motif matches, only relatively common potential phosphorylation sites could be identified. Database searches showed that homologues were restricted to Pandoraviruses.

For Cluster 12, different modelling methods were tried, since an initial AF2 model had only a mean pLDDT confidence score of 58 (Fig 18). Furthermore, the most confident AF2 model positioned only two of the conserved Cys residues suitably for disulphide bond formation, while it was considered likely (also considering the predicted subcellular localisation) that four disulphide bonds would form. The emergence of these assumed disulphide bonds offered valuable extra validation of the modelling since they are not explicitly considered in Deep Learning-based modelling. A RoseTTAFold model was more confident and connected two cysteine pairs Cys44-Cys46 and Cys40-Cys57, but the most favoured model came from OmegaFold and clearly connected three disulfide bonds - Cys31-Cys46 Cys40-Cys57 and Cys49-Cys66 - with the two unconnected Cys44 and Cys79 close enough to make a fourth bridge plausible (Fig 18).

**Figure 18.**
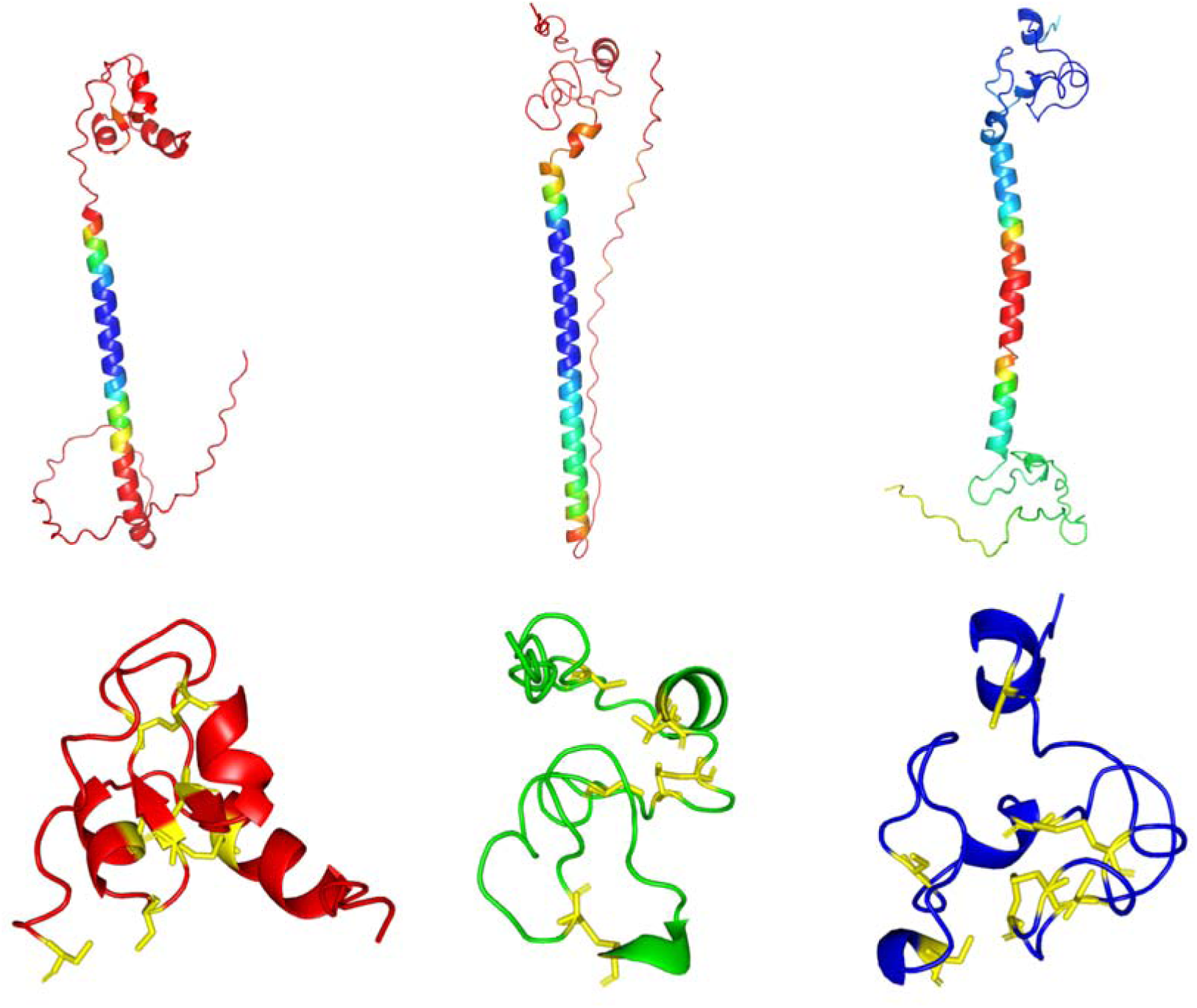
Top: S4VS46 modelled by OmegaFold (left), AF2 (centre) and RosETTAFold (right). OmegaFold and AF2 model coloured by confidence, RosETTAFold model coloured by angstroms error estimate (not using AF2 colour scheme). Bottom: Closer view of the globular extracellular region (residues 19-79 shown) of the OmegaFold (left), AF2 (centre) and RosETTAFold (right) models, with 8 conserved cysteines coloured yellow. The globular regions contain three, one and two disulphide bonds, respectively, in the three models.

Of the three models, only the globular region of the OmegaFold gave hits in the AFDB version 3 using FoldSeek that registered TM-scores above 0.6. The sequences of these were searched against the PDB using HHpred resulting in 26 unique hits with >80% probability. Ultimately, four hits strongly resembled S4VS46 in structure and disulphide connectivity: 1olz-A (residues: 467-523, connectivity: 482-499, 491-508), 1ssl-A (residues: 2-48, connectivity: 6-24, 12-47, 15-31, 27-37), 7mob-C (residues 516-554, connectivity: 529-545, 520-538, 541-551), and 5l74-A (residues: 654-702, connectivity: 662-701, 656-672, 665-681, 675-688) (Fig 19). These PDB structures with structure and disulphide connectivity were considered as potential distant homologs of Cluster 12 proteins that were not discernible by simple sequence comparisons.

**Figure 19.**
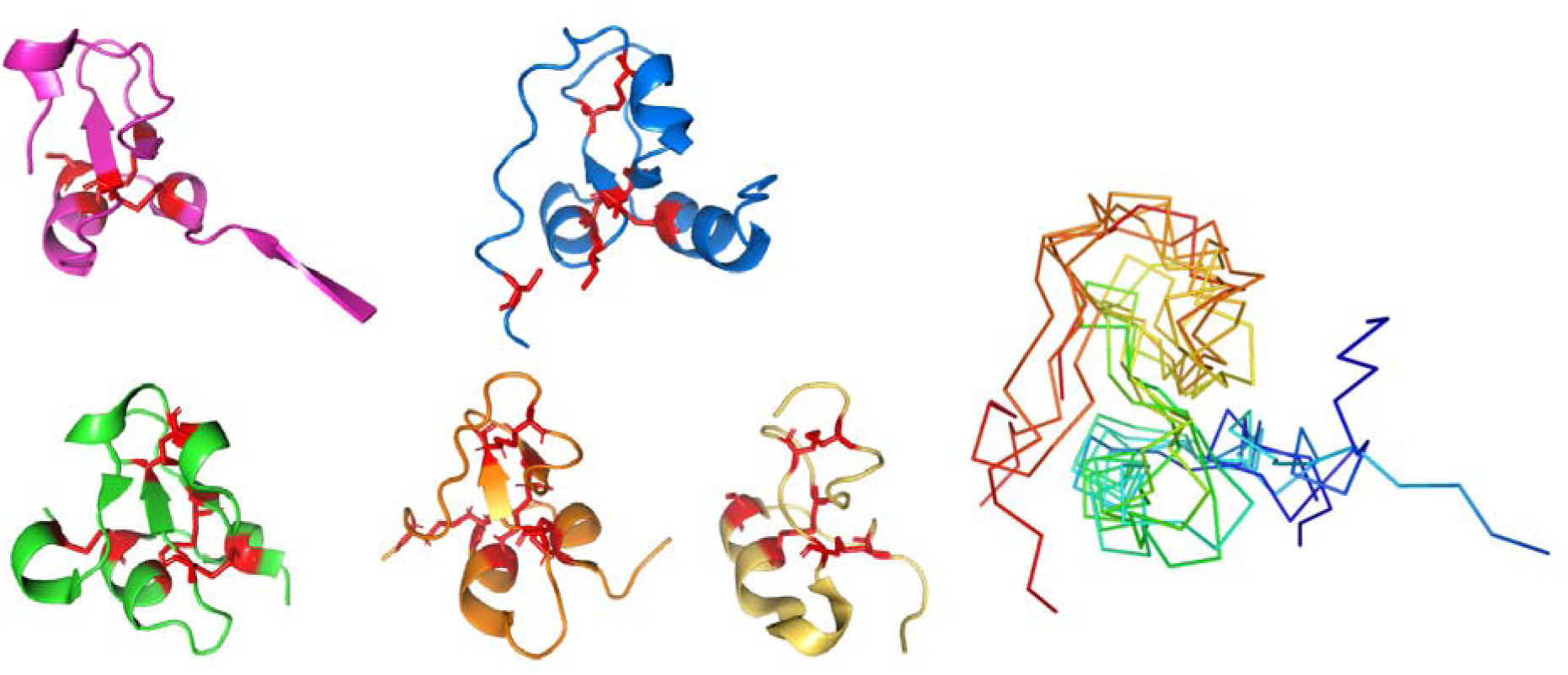
Left: Diverse PSI domain homologues with identical connectivity to S4VS46 (blue) with cysteines shown in red: 1olz-A (magenta; residues 474-523), 5l74-A (green; residues 654-702), 1ssl-A (orange; residues 2-48), 7mob-C (yellow; residues 516-554). Right: superposition of all five proteins, coloured from C terminus (blue) to N-terminus (red), showing identical topology. All proteins aligned to S4VS46 in PyMOL using cealign.

These four domains, in 5l74-A, 7mob-C, 1ssl-A and 1olz-A are found in receptors for semaphorin, Met and SEMA4D and are annotated as PSI domains in databases such as Pfam. The PSI domain functions structurally to link the stalk region and the ligand recognition domain in Met (Kozlov *et al*., 2004); (Uchikawa *et al*., 2021), and orients the ligand binding and structural domains in SEMA4D (Love *et al*., 2003). Interestingly, (Love *et al*., 2003) commented that DALI struggled to find structural matches to this cysteine knot topology, as was also seen here for S4VS46. PSI domains are present in extracellular regions of hundreds of signalling proteins, and share little sequence identity apart from six invariant and two highly conserved cysteines. A tryptophan is also conserved (Love *et al*., 2003; Kozlov *et al*., 2004), and can be identified as invariant across cluster 12 e.g. at position 48 in S4VS46. These detailed considerations suggest that Cluster 12 encodes a divergent PSI-like domain, but one divorced from its usual receptor context.

InterPro Taxonomy showed that PSI domain-containing proteins are mostly found in eukaryotes, with only 21 viral protein sequences, two from megaviruses. However, all 21 proteins were predicted to be semaphorins or Sema domain-containing proteins, and according to DeepTMHMM none contained transmembrane helices. This makes Cluster 12 proteins, being transmembrane and not fused to any other globular domains, unique amongst viruses.

GO terms attached to transmembrane proteins, like S4VS46, that contain a PSI domain are generally limited to membrane localisation or are based on additional domains not present in S4VS46. For structure-based function annotation, the OmegaFold model was submitted to STP, which gave no clear predictions, and CSM-potential, which identified two potential protein-protein interaction sites on the transmembrane helix. Consurf was limited to two iterations of HMMER against UniProt (to avoid contamination of the database search with Cys-rich but unrelated sequences) and showed the globular region as most conserved. Thus, the function of this cluster remains cryptic.

#### Cluster 13

Cluster 13 contained 13 protein sequences between 182 and 271 residues long. Sequence analysis suggested a transmembrane protein with two helices and localisation in the endoplasmic reticulum (ER). No apparent homologues were found outside of pandoraviruses. AlphaFold2 was used to predict a structure with high confidence in the transmembrane helices, moderate confidence in a small, all-β intra-lumenal domain inside the endoplasmic reticulum and lower confidence in linker regions and at the N-terminus (Fig 20a). Unsurprisingly, given the small size of the confidently folded parts, searches of structure databases failed to reveal strongly similar structures. Structure-based function using STP supported the possibility of an possible interface on the surface of the two transmembrane α-helices (Fig 20b), one of which, along with the small β-domain, is highly conserved (Fig 20c).

**Figure 20.**
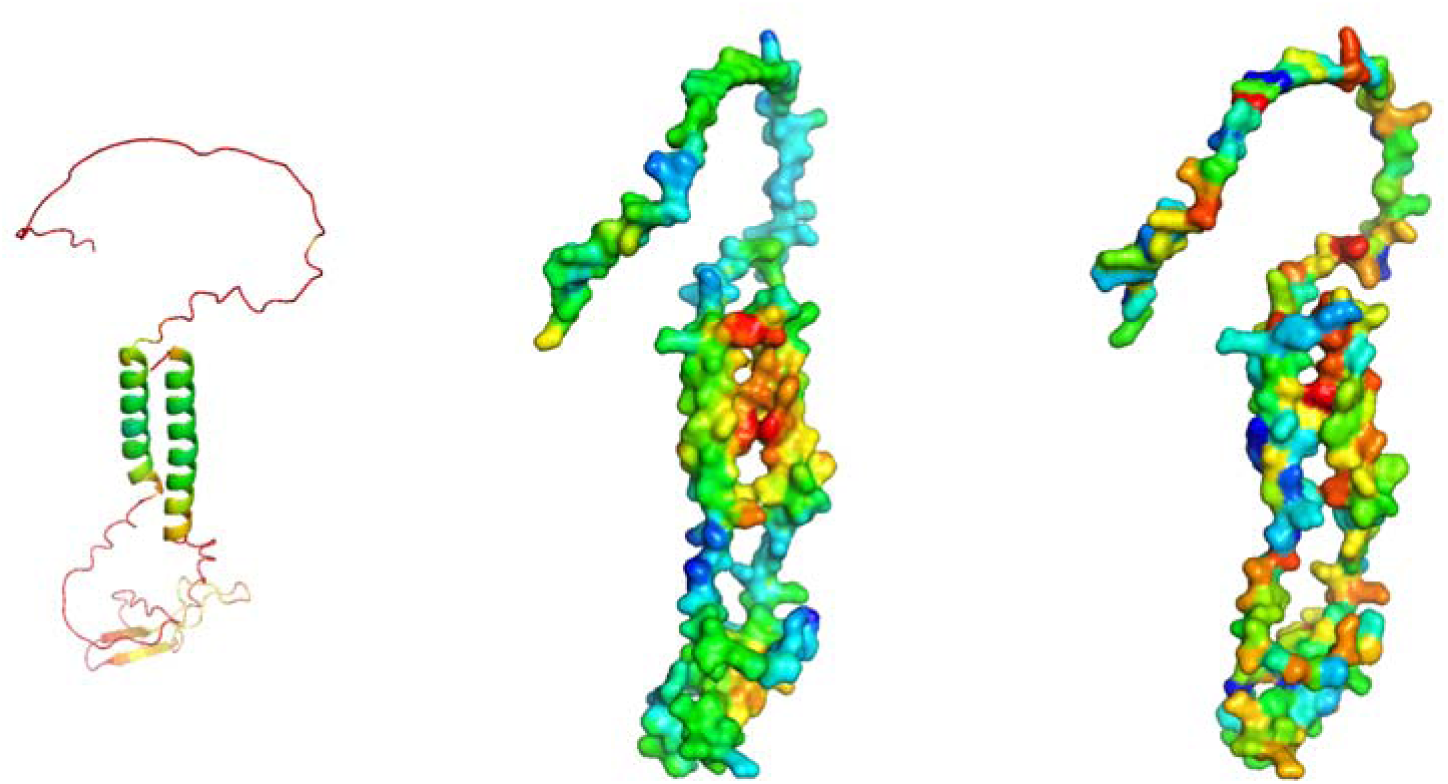
Structural analysis of S4VZF8. (a) AF2 prediction, displaying high structure prediction of the two α-helices. (b) STP analysis, potential interaction surface on the two α-helices shown in red/orange. (c) ConSurf conservation analysis where blue indicates conservation and red unconserved residues.

Proteins with two transmembrane helices can oligomerise for function, as for example is the case of the KcsA potassium channel (Zhou *et al*., 2001). To explore possible oligomeric states, the protein was modelled as a dimer, trimer and tetramer with AlphaPullDown (Yu *et al*., 2023) and the resulting quality metrics from AF2 used to infer whether modelled interfaces are confident. As a tetramer, the transmembrane portion had the most confident interface scores, although they were still poor, and produced an arrangement that, speculatively, could form a channel pore.

Ultimately, the hypothesis that cluster 13 encode tetrameric potassium channels must be considered speculative, although it is intriguing to note that the ER is known to harbour such channels and that other megaviruses, the algal chloroviruses, include several characterised two-helix potassium channels (Thiel *et al*., 2013).

#### Cluster 14

Cluster 14 had 12 sequences in total, ranging between 87 and 119 residues long. Interestingly, proteins in this cluster were either ∼90 residues or ∼110 residues long, the difference between the two being a longer C-terminal in the longer members. A0A2U7U965, a longer member with 116 residues, was chosen as a representative. Database searches showed obvious homologues only in Pandoraviruses, with HHpred not identifying any more distant relationships with PDB structures or Pfam families. DeepLoc predicted extracellular localisationand other sequence-based predictions returned negative results (Table 1).

Alphafold2 was used to model the cluster representative (Fig 21). The model displayed a beta sandwich structure with a high confidence. Interestingly, conserved Cys residues, numbers 10 and 59 in the representative sequences, are found close together in the model (although Cys 10 is found in the low-confidence N-terminal region) and could potentially form a disulphide bond. Such a bond would be consistent with the predicted extracellular localisation. Interestingly, these two cysteines are present or absent in a coordinated fashion in homologues supporting the existence of the disulphide bond.

**Figure 21.**
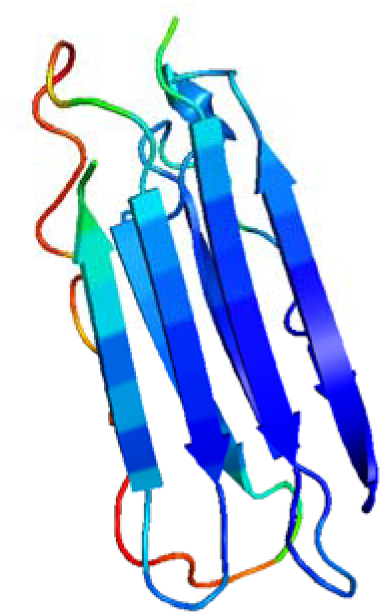
AF2 model of Cluster 14 representative A0A2U7U965 coloured by confidence scores (blue high, red low). N-terminal residues 1-16 with pLDDT confidence values of < 50 are not shown.

Structural neighbours in the PDB were then sought with DALI. A large number of significant hits with diverse functions were returned (Table 2). Many of these functions were considered unlikely: for example, the match (Z-score = 8) to ostreolysin A, a sphingomyelin-cholesterol complex-binding protein from *Pleurotus ostreatus* (PDB: 6MYI; (Endapally *et al*., 2019)), conflicts with the understanding that sphingomyelin is not known to be found in Acanthamoebae (ref). Similarly unlikely is the match (Z-score =7.8) to the SPH domain in 6G7G. That structure is a member of the SPH (self-incompatibility protein homologue) family whose binding to pollen receptors is involved in avoidance of self-fertilisation in plants (Rajasekar *et al*., 2019). Ultimately, fold-based function prediction is simply less valuable for structures with common folds such as the β-sandwich adopted here. Instead, bottom-up structure-based function annotation was attempted. However, neither electrostatic or STP analysis revealed any likely binding sites.

## Conclusions

Alphafold 2 has changed the landscape of protein structural bioinformatics, providing not only coordinates of often unprecedented accuracy, but also excellent confident estimates that enable researchers to know when to trust the predictions. It is therefore an invaluable tool to mine the dark proteome i.e. proteins whose relationships to known families or structures are not close enough to allow for confident functional assignment. The AlphaFold Database includes models for a large proportion of the UniProt protein sequence database, but virus proteins are currently largely absent (Varadi *et al*., 2022).

This work focused on Pandoraviruses since they contain a conspicuously large proportion of ‘hypothetical proteins’ hinting at much interesting biology still to be discovered. By targeting families of hypothetical proteins, rather than individual sequences, we aimed to annotate multiple proteins on the basis of detailed scrutiny of representative sequences. Indeed, in several cases we provide strong functional predictions for newly described protein families present in multiple copies in diverse Pandoraviruses. To our knowledge, cluster 5 represents the first membrane transporters known to be encoded by these viruses: charge analysis suggests a negatively charged substrate but the identity of the transported ion or molecule must await experimental study. Cluster 2 is very likely to encode regulatory beta subunits of a calcium-activated potassium channel, yet we could not find the predicted alpha subunit partner in either virus or known hosts: this may hint at an unsuspected ability of these viruses to infect further hosts. Cluster 7 contains class II nucleotidyltransferases likely of relaxed nucleotide specificity that may be involved in maturation of the pandoravirus-encoded tRNAs. The BTB domain, also found in cluster 7, resembles that found in proteins that interact with Cul3, suggesting that - like diverse other viral proteins - it may be involved in manipulation of the host cell’s ubiquitin-dependent pathways, to the benefit of the Pandoravirus.

AF2 is known to be able to accurately predict novel folds distinct from anything in its training set (Jumper *et al*., 2021) making it useful for computational characterisation of structural novelty until the PDB provides a complete picture of protein foldspace (Durairaj *et al*., 2023; Hernandez *et al*., 2023). For example, the work of Bordin et al in the context of the CATH database revealed a number of new structural superfamilies, even in an analysis of an earlier AlphaFold Database focussed on model organism proteomes (Bordin *et al*., 2023).

Limitations encountered in the current work include fold matches for new predictions (such as those for clusters 4 and 14) which, while significant, were to functionally promiscuous families providing no strong steer as to likely function. The relative paucity of Pandoravirus sequences also proved problematic since it reduced the signal available from sequence mapping onto structure, generally a reliable and powerful means to locate functional sites (Rigden, 2017). A greater number of available homologues (which is naturally expected with time) would also be expected to have a favourable impact on model quality, reducing or eliminating the model regions eg in cluster 7 that were of lower confidence but which, from a relative richness of predicted regular secondary structure for example, were likely to be folded rather than intrinsically disordered. Finally, while new structure-based function annotation methods continue to emerge (Cagiada *et al*., 2023; Krapp *et al*., 2023), it is currently frustrating that older generation yet still potentially valuable methods like STP (Mehio *et al*., 2010; Krapp *et al*., 2023) have not yet been benchmarked on AF2 structures.

Despite these issues, this work annotated intriguing functions onto several newly defined families and revealed novel folds in others: it should encourage similar work to employ the latest modelling methods to characterise *in silico* the dark matter of other viruses and cellular organisms.

## Data availability

Models presented in this manuscript are available via FigShare at https://doi.org/10.6084/m9.figshare.24701700.v2

