## Supplementary Material for "Deep Learning-based structural and functional annotation of Pandoravirus hypothetical proteins"

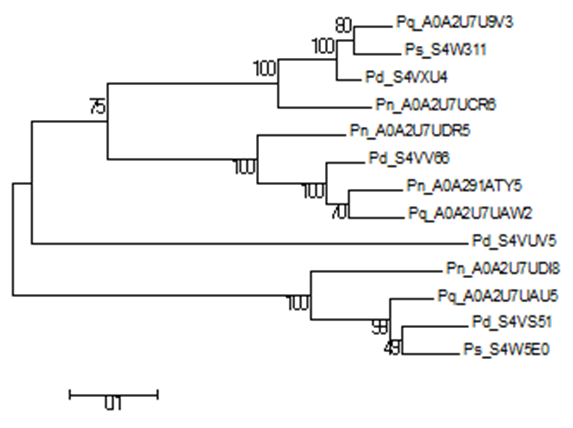

2

3

1

**Supp Figure 1.** Molecular Phylogenetic analysis of Cluster 5 by Maximum Likelihood method

The evolutionary history was inferred by using the Maximum Likelihood method based on the JTT matrix-based model [1]. The tree with the highest log likelihood (-5460.3488) is shown. Bootstrapping of 1000 replications was carried out. The percentage of trees in which the associated taxa clustered together is shown next to the branches. Evolutionary analyses were conducted in MEGA6 [2].

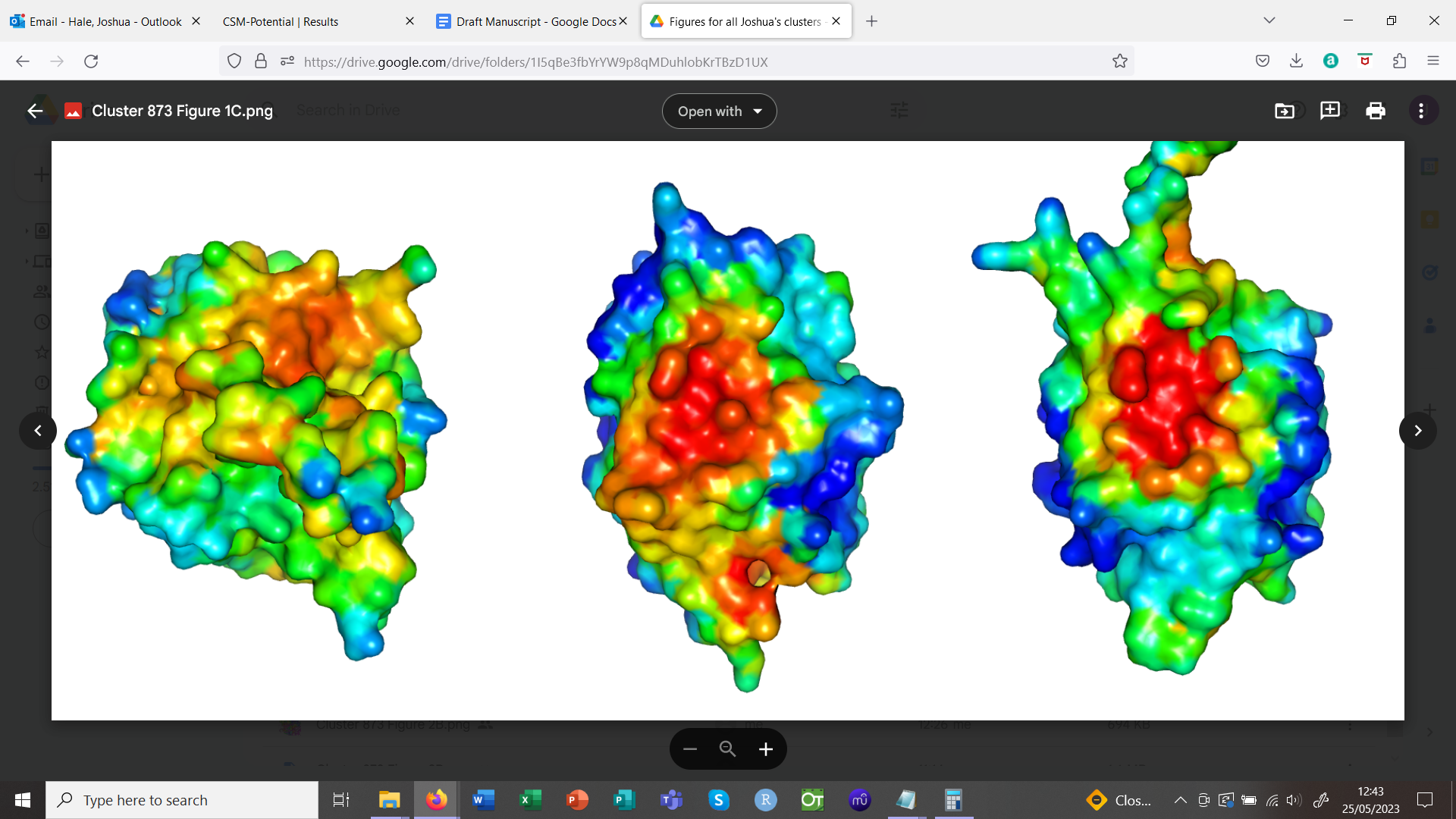

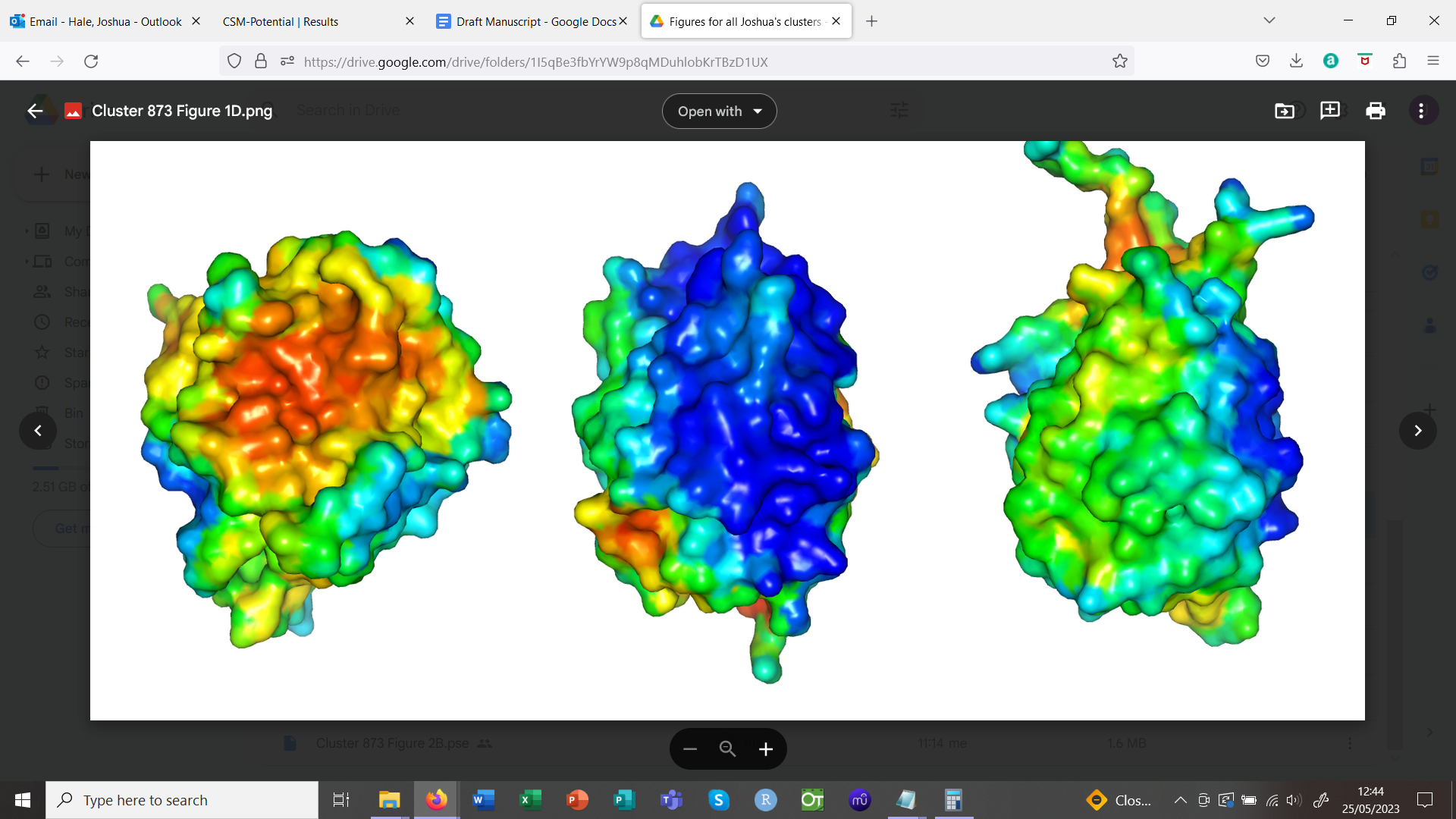

**Supp FIgure 2:** CSM-potential protein-protein interaction predictions for aligned cluster 4 representative A0A2U7U8Z7 (left; front above, rear below), *Arabidopsis thaliana* SCDF2L1 (3mal_B; centre) and an AF2 model of *Mus musculus* SCDF2L1 (right). Figure coloured red for strong prediction through to blue for no predicted interface.

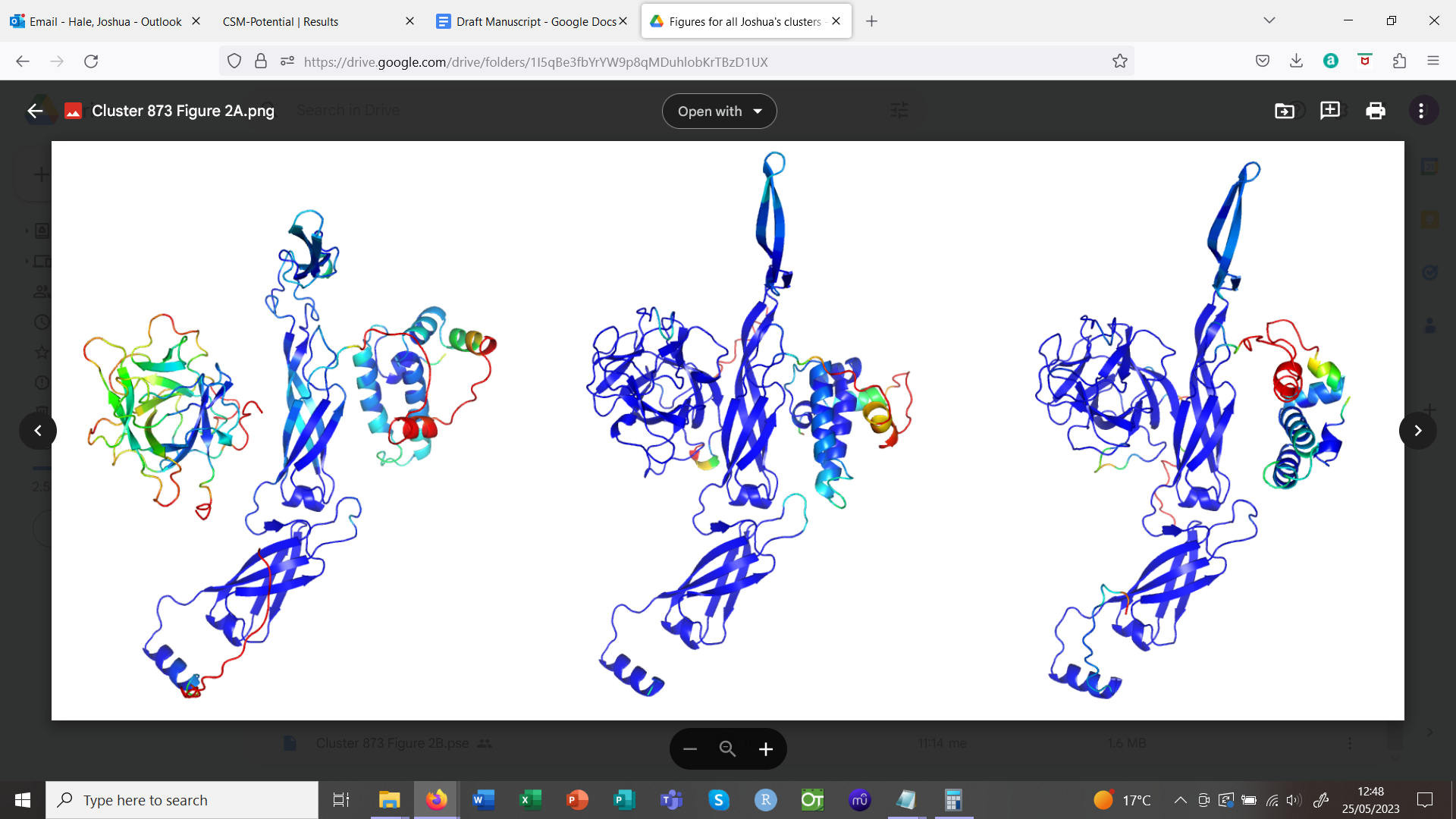

**Supp Figure 3:** Top: comparison of AF2 heterodimer models (coloured by confidence): cluster 4 representative A0A2U7U8Z7 with putative amoebal ERdj3 chaperone L8H8X3 (left), 3mal_B with *Arabidopsis thaliana* ERdj3 (centre) and *Mus musculus* SCDF2L1 with *M. musculus* ERdj3 (right). All heterodimers aligned to A0A2U7U8Z7/L8H8X3 in PyMOL using cealign (Shindyalov and Bourne, 1998).

**Supp Table 1.** Characteristic conserved residues in the three clades of the Cluster 5 phylogenetic analysis

| **Residue in Cluster 5 reference sequence** | **Clade 1*** | **Clade 2** | **Clade 3** |
| --- | --- | --- | --- |
| S64 | A | T | S |
| G78 | N | T | G |
| L79 | F | Y | L |
| L140 | F | Y | L |
| P141 | L | S | P |
| A175 | F | L | A |
| A186 | G | D | A |
| A198 | L | I | A |
| L215 | I | F | L |
| A220 | K | G | A |
| V268 | A | Q | V |
| S272 | V | M | S |
| G331 | A | S | G |
| A334 | G | V | A |
| G385 | N | T | G |
| F422 | L | V | F |
| P447 | A | S | P |
| V460 | T | A | V |
| A469 | V | T | A |
| P475 | V | M | P |

*Sequence A0A2U7UCR6 excluded from the analysis since it appears partial

**Supp Table 2:** DALI hit representatives (C4A = Cluster 4 alignment by MAFFT)

| **Pfam Domain** | **Representative Hit (Z-score)** | **Key features and presence/absence in A0A2U7U8Z7 and Cluster 4 Alignment.** | **References** |
| --- | --- | --- | --- |
| PF02815: MIR domain | 3uj4-A (15.7) | 3uj4-A binding site composed of conserved Arg residue and other domain interfaces, absent in A0A2U78Z7.  Conserved Pfam Trp and Leu residues absent in C4A. | (Yoshikawa *et al.*, 1996; Lee *et al.*, 2016; Bai *et al.*, 2019; Chiapparino *et al.*, 2020) |
| PF08709: Inositol 1,4,5-trisphosphate/ryanodine receptor | 3im5-B (15.0) | 3im5-B irrelevant function: interdomain interactions within the ryanodine receptor 2. | (Lobo and Van Petegem, 2009; Kimlicka *et al.*, 2013) |
| PF07468: Agglutinin domain | 1jly-B (15.0) | Conserved T-antigen disaccharide binding residues (Asn, His, Tyr and Trp) and similar binding site in 1jly-B absent in both C4A and Consurf. | (Transue *et al.*, 1997) |
| PF07951: Clostridium neurotoxin, C-terminal receptor binding | 3n7j-A (14.7) | Conserved sialic acid and ganglioside binding residues (Thr, Asp, Lys, Arg, Tyr and Val) and similar binding sites in 3n7j-A absent in both C4A and Consurf. | (Karalewitz *et al.*, 2010); (Strotmeier *et al.*, 2010) |
| PF08470: Nontoxic nonhaemagglutinin C-terminal | 3v0b-B (14.6) | 3v0b-B irrelevant function: a domain of the NTNHA protein involved in interdomain interactions and protein-protein interactions with *Clostridium botulinum* neurotoxin. | (Gu *et al.*, 2012) |
| PF00197: Trypsin and protease inhibitor | 5hpz-B (13.9) | 5hpz-B irrelevant function: conserved chlorophyll-binding Pro residues absent in C4A and *Acanthamoeba spp.* not photosynthetic.  Typically conserved Pfam Arg, Asn or Lys residues and two disulfide bridges for the protease inhibitory loop absent in C4A. Whilst three disulfide bridges are present in A0A2U7U8Z7, only one of these is between Cys residues conserved in the C4A. | (Bednarczyk *et al.*, 2016; Bendre, Ramasamy and Suresh, 2018) |
| PF14200: Ricin-type beta-trefoil lectin domain-like | 3win-B (13.8) | Conserved Pfam QxW motif in absent in C4A.  A0A2U7U8Z7 matches the protein-protein interaction region, and not the carbohydrate binding region, of 3win-B. | (Nakamura *et al.*, 2008, 2011; Rotskaya *et al.*, 2021; Xu *et al.*, 2022) |
